# Voltage imaging reveals circuit computations in the raphe underlying serotonin-mediated motor vigor learning

**DOI:** 10.1101/2024.09.15.613083

**Authors:** Takashi Kawashima, Ziqiang Wei, Ravid Haruvi, Inbal Shainer, Sujatha Narayan, Herwig Baier, Misha B. Ahrens

## Abstract

As animals adapt to new situations, neuromodulation is a potent way to alter behavior, yet mechanisms by which neuromodulatory nuclei compute during behavior are underexplored. The serotonergic raphe supports motor learning in larval zebrafish by visually detecting distance traveled during swims, encoding action effectiveness, and modulating motor vigor. We found that swimming opens a gate for visual input to cause spiking in serotonergic neurons, enabling encoding of action outcomes and filtering out learning-irrelevant visual signals. Using light-sheet microscopy, voltage sensors, and neurotransmitter/modulator sensors, we tracked millisecond-timescale neuronal input-output computations during behavior. Swim commands initially inhibited serotonergic neurons via GABA, closing the gate to spiking. Immediately after, the gate briefly opened: voltage increased consistent with post-inhibitory rebound, allowing swim-induced visual motion to evoke firing through glutamate, triggering serotonin secretion and modulating motor vigor. Ablating GABAergic neurons impaired raphe coding and motor learning. Thus, serotonergic neuromodulation arises from action-outcome coincidence detection within the raphe, suggesting the existence of similarly fast and precise circuit computations across neuromodulatory nuclei.

## Introduction

Neuromodulation alters behavior, allowing animals to adapt to changes in their environment, respond to salient interactions with other animals, and adjust to unexpected signals from body organs^1^. These behavioral adjustments occur as secreted neuromodulators bind to receptors on neurons and astrocytes, shaping circuit dynamics^2,3^. While much research has focused on understanding the effects of neuromodulation on circuit dynamics and behavior, as well as the computational interpretation of variables encoded in the spike patterns of neuromodulatory cells^4^, there remains a gap in our knowledge of how internal circuits, within neuromodulatory nuclei, process information during behavior. To trigger behavioral adjustments, how do cells within these nuclei interact with one another to transform input signals from other brain regions into precisely timed neurotransmitter release?

A significant behavioral change occurs when animals adjust their motor commands to meet the demands of dynamic environments. For example, when walking against a strong wind, humans lean forward and apply more force to maintain their pace. Similarly, fish swimming in currents must adapt their swim vigor to the speed of water; if low temperature weakens their muscle contractions or changing water viscosity alters the effectiveness of their tail in propelling their body forward (i.e., *action effectiveness*), they modify swim vigor accordingly. Such adjustments rely on sensory-motor computations that evaluate sensory outcomes of motor actions and adjust future motor commands. Neural mechanisms of motor learning are distributed across the brain^5–7^, including neuromodulatory systems^8–10^. The vertebrate serotonergic system modulates motor learning at various levels, from enhancing sensory perception to directly suppressing spinal cord outputs^11^. Such modulation can have differential effects depending on behavioral context^12,13^. In zebrafish, serotonergic neurons in the dorsal raphe nucleus (DRN) drive motor vigor learning. Serotonergic neurons continuously evaluate the sensory outcomes of their swimming, such as the distance traveled per swim bout, update an internal estimate of action effectiveness in memory represented by spike rates, and modulate future motor vigor by releasing serotonin^14^. Action effectiveness must be computed from motor output and motor-triggered visual feedback, and stored for future modulation of motor vigor for many seconds. Yet, it remains unknown how these motor-sensory computations occur via network and cellular interactions, specifically within the serotonergic system. Such insights are necessary to understand how the vertebrate serotonergic system integrates multiple streams of sensory and motor information to control behavior ^4,15–20^.

Larval zebrafish learn and adapt motor vigor to changes in action effectiveness within a dynamic environment. The same swim bouts may not consistently result in the same traveled distance; fish thus need to monitor the distance traveled per swim and adjust their swim vigor accordingly^21,22^. If swim commands cause larger-than-intended body displacements, swim vigor decreases. Conversely, if swim commands cause little displacement, swim vigor increases. While passive displacement of the fish’s location (e.g. resulting from sudden water currents) can impact swimming^23^, only self-produced displacements should drive motor learning. More generally, learning action effectiveness requires animals to use temporal correlations between actions and feedback to accurately determine the resulting outcomes to be used in motor learning.

Here, we investigated the input-output transformations, and the underlying millisecond-timescale mechanisms, of DRN serotonergic neurons during motor vigor learning in larval zebrafish. A visual virtual reality environment allowed us to control action effectiveness explicitly while imaging a suite of genetically-encoded voltage and neurotransmitter indicators in serotonergic neurons using light-sheet microscopy. The temporal pattern of axonal serotonin release was consistent with a theoretical model for integrating action effectiveness over time. During swims, serotonergic neurons’ membrane potential and spiking activity is momentarily suppressed but then recovered from inhibition to encode visual displacement, i.e., action effectiveness. Direct observation of excitatory and inhibitory inputs and membrane voltage, together with theoretical simulation and precise perturbations, suggests a model in which the encoding of action effectiveness results from the synergistic interaction of neuronal biophysics with incoming sensory and motor signals onto serotonergic neurons. When visual input arrives immediately after a swim command, visually-driven excitation is added to voltage increases that result from swim-evoked post-inhibitory rebound^24–27^ to cause spiking in serotonergic neurons that encodes the visual stimulus. Such spiking does not occur when visual input arrives outside of a specific time window after a motor command due to the absence of voltage rebound, implementing a cellular coincidence detection mechanism between actions and outcomes. Generalized forms of this mechanism of action-outcome computation may operate in the vertebrate serotonergic system across behavioral contexts.

## Results

### Zebrafish serotonergic system learns action effectiveness to adapt motor vigor

To measure how the serotonergic system computes action effectiveness for each swim event in zebrafish, we first measured axonal serotonin (5-HT) release from the dorsal raphe nucleus while the fish performed a motor vigor learning task in a virtual reality (VR) environment (**Fig. 1a**). In this virtual visual environment (**Fig. 1b**), a paralyzed head-fixed fish swam against a virtual water current, which is simulated by constant forward motion of the visual environment projected on a screen beneath the fish. Their swim signals from the spinal cord were detected in real-time through a pair of electrodes connected to the skin of the tail. When a swim signal was detected, fish moved forward in VR, which was simulated as transient backward motion of the projected environment. The velocity and distance of swimming were proportional to the amplitude of detected swim signals, scaled by motosensory gain (Gms), which represents action effectiveness and can be arbitrarily manipulated to induce motor vigor learning^21,22^. Fish adapted the amplitudes of swimming signals in response to changes in Gms in a compensatory manner: low Gms induced progressively stronger swimming, and high Gms induced progressively weaker swimming (**Fig. 1b**).

**Figure 1.**
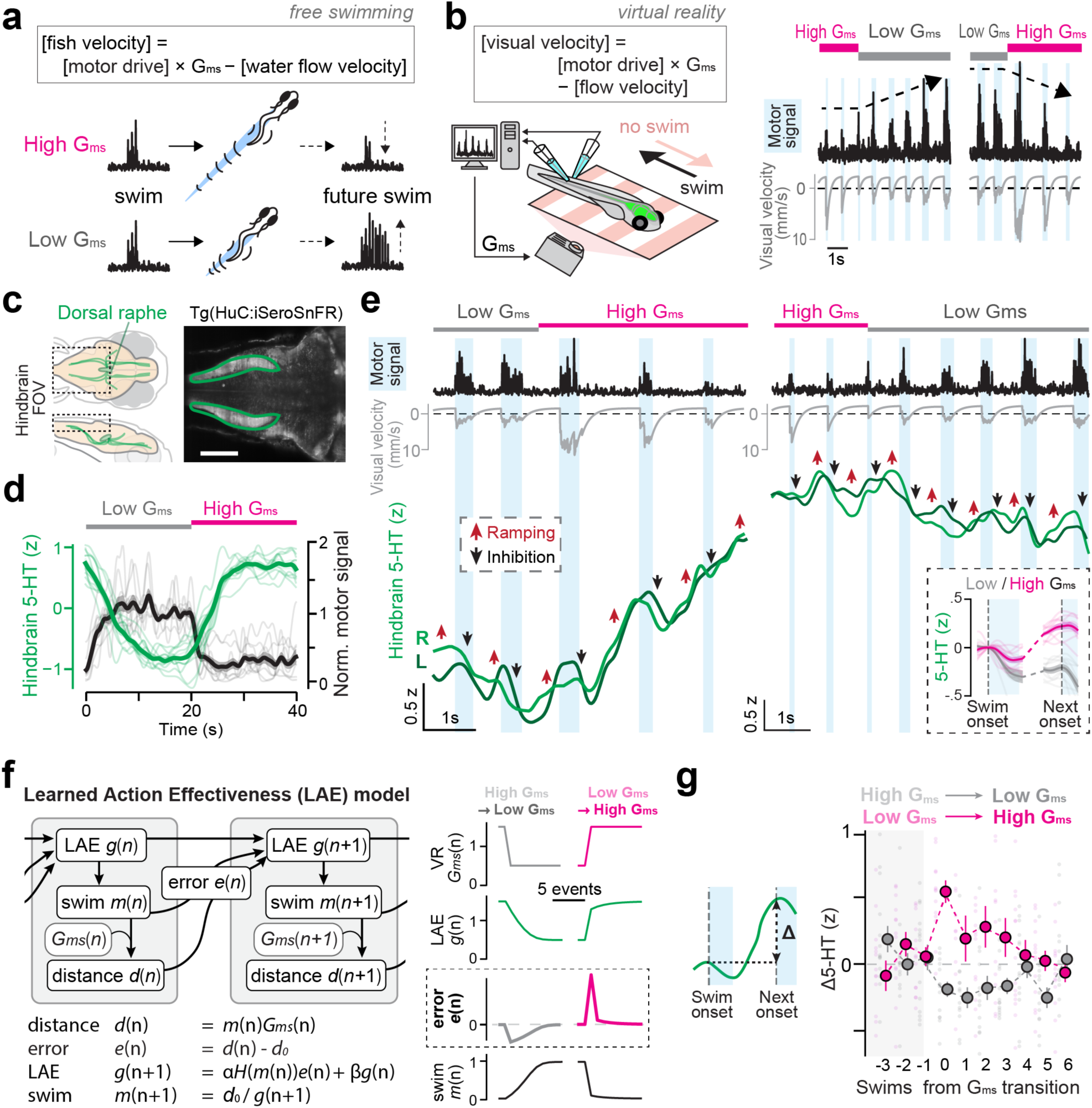
Serotonin release pattern during motor vigor learning reflect learning of action effectiveness. **(a)** Schematic of motor adaptation in response to changes to a high or low level of action effectiveness (G_ms_). Animals need to exert more effort to travel the same distance in low G_ms_ than in high G_ms_. To uphold a constant traveled distance per action, animals adapt by intensifying their effort per action in low G_ms_ and reducing it in high G_ms_. **(b)** *Left,* experimental setup with virtual reality environment. In this setup, action effectiveness was controlled by a computer program via parameter G_ms_. *Right,* swimming and visual feedback during low and high G_ms_. **(c)** Schematic of the zebrafish raphe serotonergic system (left) and an image of HuC:iSeroSnFR fish with regions of interest in the hindbrain neuropil downstream of the serotonergic raphe (green, right). Scale bar, 100 µm. **(d)** Larval zebrafish adapt within ∼10 seconds to changes in action effectiveness, with increases in motor vigor during low G_ms_ and decreases in vigor during high G_ms_ (black). Serotonin release in the hindbrain (z-scored fluorescence, green) showed opposite temporal patterns. N=9 fish. **(e)** Serotonin signals (z-scored fluorescence), shown swim-bout-by-swim-bout in ROIs of (c), exhibit initial inhibition of the 5-HT signal during swimming (black arrow), followed by an increase immediately after (red arrow). At high G_ms_, the increases are stronger and more sustained, leading to ramping activity. L and R indicate serotonin release timeseries from the hindbrain neuropil of the left and right side, respectively. Single trial data. *Inset:* serotonin release pattern (z-scored fluorescence) averaged across swim bouts showing increased 5-HT release following swim bouts in high G_ms_ compared to those in low G_ms_. **(f)** Schematic (top) and algorithmic implementation (bottom) of *Learned Action Effectiveness (LAE)* model. Parameters α, learning rate; β, retention factor; d_0_, a constant representing desired distance to travel per swim. Our model was updated at swim events, n. *Right:* Simulations of an *LAE* model during 10 swim events (equivalent to ∼10 seconds for real fish) at the transition from low G_ms_ to high G_ms_, and the transition from high G_ms_ to low G_ms_ during motor vigor learning. **(g)** Swim-by-swim increase of serotonin release (Δ5-HT; z-scored fluorescence) at the transitions of G_ms_ in our experiments qualitatively matches the error term in the *LAE* model. N = 9 fish.

We examined the temporal pattern of serotonin release during this motor learning task by imaging serotonin release from axonal projections of DRN serotonergic neurons in the hindbrain using a genetically-encoded indicator iSeroSnFR (**Fig. 1c**). This low-affinity serotonin indicator has fast kinematics (τdecay = 150 ms in cultured neurons) and is suited for capturing temporal dynamics of serotonin release during swimming at sub-second timescales^28^. In a motor adaptation task where Gms transitioned between low and high values every 20 seconds, fish adapted and equilibrated their swim vigor over a period of ∼10 s after the transition (**Fig. 1d**). Serotonin release in the hindbrain was inversely related to swim vigor on slow timescales. Serotonin levels increased when the fish attenuated their swim vigor and decreased as fish boosted their swim vigor (**Fig. 1d**). At a finer timescale, serotonin level showed a transient decrease during every swim bout, followed by a rise during inter-swim intervals (**Fig. 1e**). Despite the decrease during swims, post-swim 5-HT rises more at high Gms, leading to a gradual increase in 5-HT levels; at low Gms the post-swim increase was small, leading to an overall decrease in 5-HT levels (**Fig. 1e, Fig. S1A**) indicating that 5-HT levels could track an internal estimate of action effectiveness. This change of serotonin levels after swimming is predictive of the vigor of the next swim bout (**Fig. S1b**), suggesting that the internal estimate of action effectiveness drives motor adaptation in zebrafish.

We formulated a learning model, called the Learned Action Effectiveness (LAE) model, to explain this form of motor learning (**Fig. 1f**) and compared its predictions with the temporal dynamics of serotonin release. In this model, with the variable n indexing swim bout number, the actual travel distance after a swim bout *d*(n), which is estimated from visual input during and right after a swim event *m*(n), is compared against a desired travel distance (constant *d_0_*), and the error *e*(n) between *d*(n) and *d_0_* is used to update the learned estimate of action effectiveness *g*(n+1). Fish generate the next swim event with vigor *m*(n+1) based on the desired travel distance *d0* and estimated action effectiveness *g*(n+1). This *g*(n+1) is only updated when there is a motor action, captured by Heaviside function *H* which equals 1 when its argument is positive and 0 otherwise, and decays with time with a retention coefficient *β*. We included these two factors in the model to explain our previous observation of sensory gating, in which sensory responses only occur during a short time window following a motor command, and the slow decay of tonic firing in DRN serotonergic neurons during the short-term motor learning task^14^. We simulated the model during transitions between high and low Gms (**Fig. 1f**) and compared the temporal dynamics of learned action effectiveness *g*(n) and error signal *e*(n) to the temporal dynamics of serotonin release. The dynamics of *g*(n) in the model and serotonin release in the experiment showed similar patterns at slower timescales (**Fig. 1f** compared to **Fig. 1d**). Notably, the error signal *e*(n) qualitatively mimicked the change of serotonin levels between swim bouts (Δ5-HT) after changes in Gms (**Fig. 1g** compared to **Fig. 1f**). Such similarity could not be obtained by models that do not include error computation (**Fig. S1c**). These results indicate that temporal dynamics of serotonin release convey the internal estimate of learned action effectiveness during motor learning.

We also investigated whether the LAE model can predict how experimental loss-of-function of the serotonergic system affects short-term motor learning (**Fig. S1d**). The loss-of-function of the serotonergic system can be simulated in LAE through a low retention coefficient *β*, and this model predicts less suppression of motor vigor during high Gms training and no effect of training durations on the swim patterns after a delay period (**Fig. S1e**). This prediction qualitatively matched the behavioral effects of the chemogenetic ablation of DRN *tph2*+ serotonergic neurons (**Fig. S1f**). Impaired suppression of motor vigor is also consistent with natural motor adaptation in freely swimming fish^29^. These consistencies between simulated and experimental behavior suggest that the zebrafish DRN encodes learned action effectiveness to modulate future actions, and motivated us to unravel the circuit and cellular mechanisms of such neural computations in the DRN.

### Voltage imaging of membrane potential and spiking activity reveals single-cell and population neural dynamics of DRN serotonergic neurons

To understand neural computations underlying the temporal patterns of serotonin release, we used a chemigenetic voltage indicator, Voltron^30^ to record membrane voltage signals from populations of raphe serotonergic neurons during motor vigor learning (**Fig 2)**. Unlike calcium imaging which has limited temporal resolution and does not report inhibition by principle^31^, voltage imaging allows us to track changes in membrane potential and spiking activity at the millisecond timescale. We used a transgenic zebrafish that expresses Voltron sparsely in *tph2*+ neurons in the DRN (*Tg(tph2:Gal4; UAS:Voltron)*), and performed voltage imaging of 10-20 neurons simultaneously at a speed of 300 Hz (**Fig. 2a**); we can thus directly determine the spiking dynamics per swim bout at millisecond timescales, and relate them to 5-HT release, visual input, and behavior. We created a custom motion correction, denoising, cell segmentation, and ΔF/F extraction algorithm to analyze membrane potential and spiking dynamics in individual neurons (**Fig. 2b; Movie S1**). We created a deep neural network for detecting spiking signals, which was trained using datasets of simultaneous voltage imaging and juxtacellular electrophysiology in zebrafish^30^. The trained network recognized electrically-recorded spikes with high accuracy (**Fig. 2c, S2A**). With these analysis pipelines, we were able to extract membrane potential dynamics and spiking activity simultaneously from ∼15 DRN neurons during the motor vigor learning (**Fig. 2d**). We leveraged this ability to record voltage signals from the serotonergic raphe to investigate how these neurons combine motor and sensory signals and enable motor learning.

**Figure 2.**
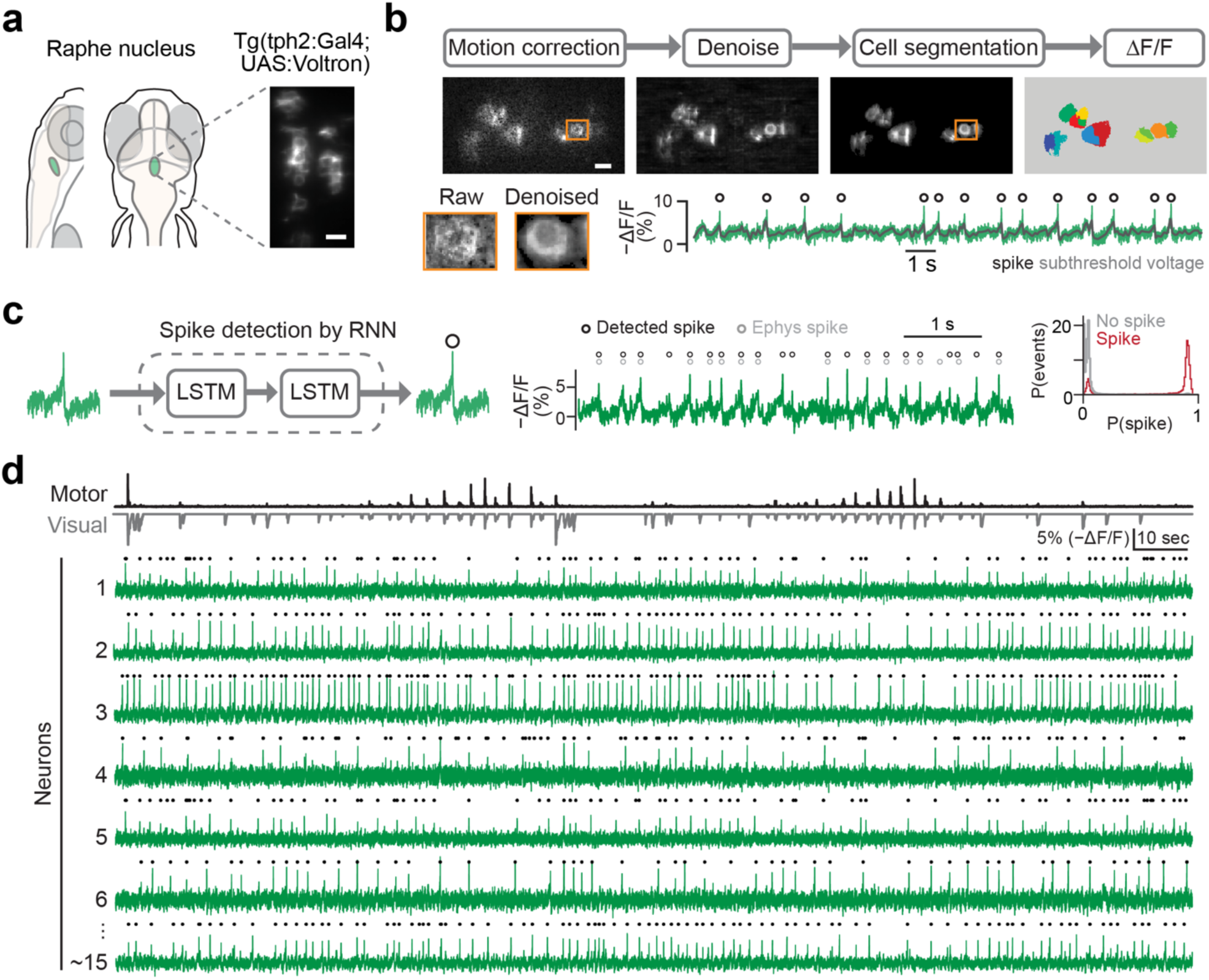
Voltage imaging in the serotonergic raphe during motor vigor learning. **(a)** Schematic of the anatomical location of the serotonergic raphe and the expression pattern of Voltron indicator in *tph2*+ neurons in transgenic zebrafish *Tg(tph2:Gal4; UAS:Voltron)*. We used JF525 dye and imaged neurons at the speed of 300 Hz. Scale bar, 10 µm. **(b)** *Top*, schematic of pre-processing pipeline: motion correction, denoising, cell segmentation, and ΔF/F calculation for individual neurons (see Methods). Scale bar, 10 µm. *Bottom*, the effects of denoising for a representative neuron marked in the top panel on the image pixel weights (left) and the estimation of subthreshold response (right) for a representative neuron marked in the top panel. **(c)** Spike-detection artificial neural network, validated with loose-patch electrical recordings. The network determines whether a spike event exists in the middle of a time series and returns a probability of a detected spike, i.e. P(spike); a spike event was assigned if P(spike)>0.5. *Left*, schematic of the recurrent neural network. *Center*, spike detection example during voltage imaging (black) and simultaneous electrophysiology (gray). *Right*, spike detection performance; x-axis, the probability of a detected spike in a time series, P(spike); y-axis, the distribution of P(spike) categorized by the ground-truth data of simultaneous voltage imaging and electrophysiology. Performances on time series with (red) and without (gray) ground-truth spike events are shown. **(d)** Voltage imaging of serotonergic neurons during motor vigor learning. *Top*, motor signals (black) and visual feedback during swimming (gray). *Bottom*, representative traces of voltage in simultaneously recorded serotonergic neurons with detected spikes.

### Motor and sensory coding during motor vigor learning

We next examined how the activity of DRN serotonergic neurons is modulated by individual swim events during motor vigor learning (**Fig. 3a**) by analyzing swim-evoked averages of membrane potential and spiking activity. We observed the suppression of membrane potential and spike rates during swims, followed by their increase after swims in individual neurons (**Fig. 3b, S2b**), in concordance with serotonin dynamics on single swim events (**Fig. 1e**). The post-swim increase in spiking was higher during high Gms, consistent with the encoding of the action effectiveness in the LAE model (**Fig. 1f,g**). This spiking pattern held true across most of the population of DRN neurons, apart from a smaller fraction of neurons that had higher, approximately-constant firing rate during low Gms (**Fig. 3c**). To disentangle the contribution of motor and sensory inputs to the post-swim response we selected swim bouts of similar vigor in both Gms conditions and found that the post-swim firing rate reflected Gms (**Fig. 3d**) and the speed of visual feedback. The increase in firing rate was fractionally larger than the increase in membrane potential, due to the averaging of post-spike hyperpolarization seen in most DRN neurons (**Fig. 2c**). Thus, during-swim hyperpolarization results from swim events, and post-swim spiking encodes the speed of visual feedback.

**Figure 3.**
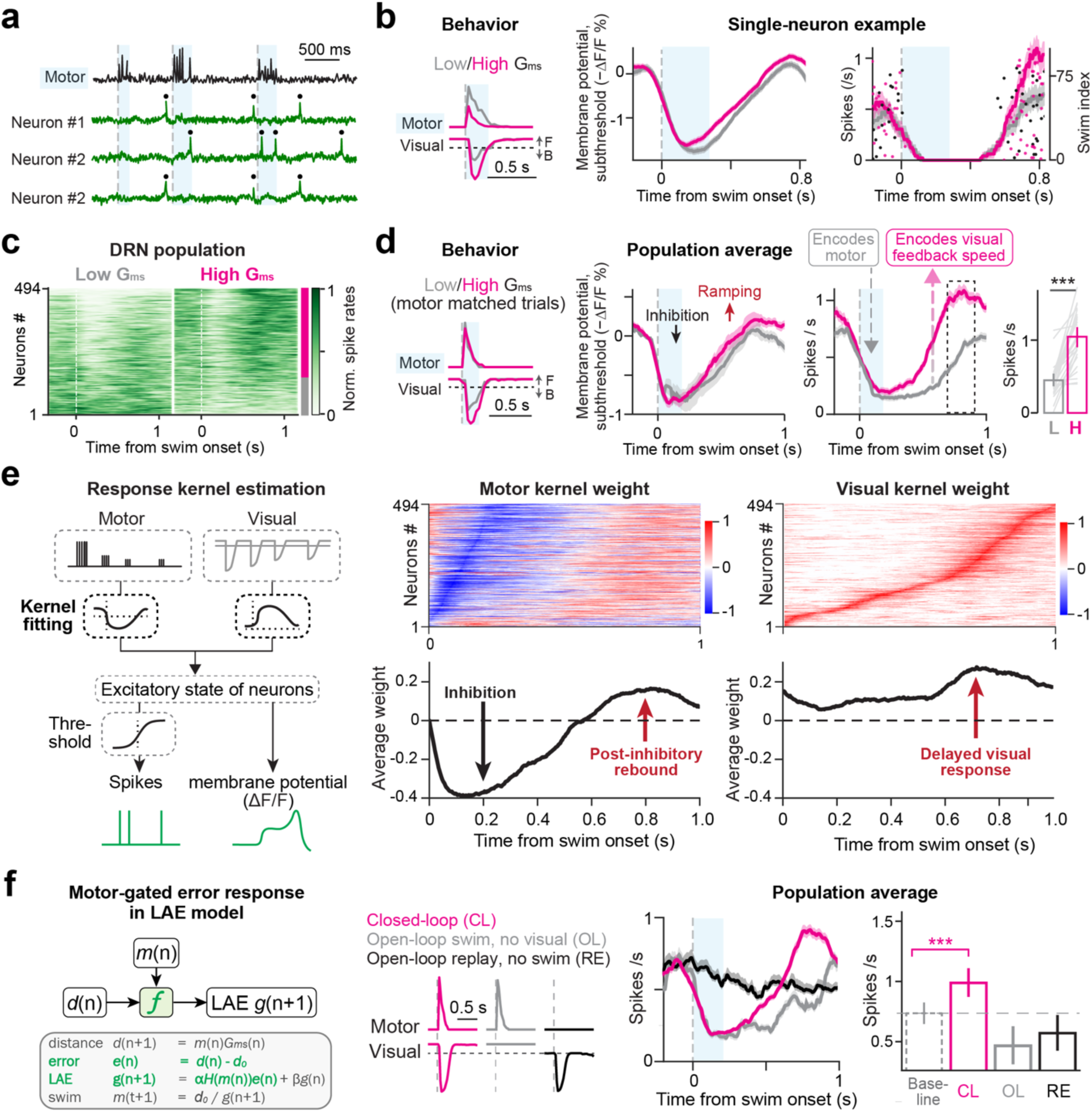
Membrane potential and spiking activity of serotonergic neurons show gain encoding and motor-gated sensory responses. **(a)** Swim signals and voltage signals from simultaneously recorded three neurons during motor vigor learning. **(b)** Changes in membrane potential and spiking dynamics of a representative neuron under different G_ms_ relative to swim bout onset. Gray, low G_ms_; magenta, high G_ms_. *Left*, swim signals and visual velocity (see Fig. 1b,e) averaged at the onset of swim events for different G_ms_ conditions; *center*, membrane potential; *right*, spike events (dots) and spike rates (lines). Shaded regions represent s.e.m. across trials. **(c)** Average spike rates of all recorded neurons at the onset of swimming for low and high G_ms_ conditions across recorded serotonergic neurons. Data of 494 cells from 21 fish are shown. The majority of neurons exhibit time-locked responses to swims in high G_ms_. High G_ms_ cells, magenta bar, N = 349; low G_ms_ cells, gray bar, N = 145. The polarity of G_ms_ of a neuron was determined by rank-sum test of spike rate after swims with a threshold of p <.05. **(d)** The velocity of visual feedback is encoded in post-swim firing rate. *Left,* average behavioral traces. We sub-selected trials to equalize motor vigor under different G_ms_ conditions. *Center,* average subthreshold and spiking responses. Dashed box represents the time window for quantifying spike rates. *Right,* spike rates after the swimming under different G_ms_. ***, p = .0007, paired signed-rank test. N = 4 fish, 29 high G_ms_ cells. Shaded areas represent s.e.m. across cells. The analysis of all swim events without the sub-selection is shown in Fig. S2b. **(e)** Fitting of a spiking kernel model (left) estimates response kernels for swimming and visual feedback in serotonergic neurons (right). Swimming triggers initial inhibition followed by post-inhibitory rebound, whereas visual feedback excites serotonergic neurons after swimming. See Methods for details. **(f)** Spiking response of serotonergic neurons to visual feedback is gated by motor actions. *Left,* gating operation is part of the LAE model presented in Fig. 1f. *Center left*, we compared three types of trials: swimming with closed-loop high-G_ms_ feedback (magenta), swim only without visual feedback (gray), visual input only (black, a replay of high-G_ms_ visual feedback). We sub-selected swim events to equalize swim vigor between closed-loop and swim only events for this analysis. *Center right,* modulation of spike rates during three trial types. Only closed-loop swims with high-G_ms_ visual feedback show post-swim spike rate increases. Subthreshold responses are shown in Fig. S2d. *Right:* changes of spike rates after swimming under tested conditions. ***, p < .0001 by paired signed-rank test of pre-swim vs post-swim spike rates. N = 7 fish, 51 high G_ms_ cells.

To formalize these findings we fit a spiking model based on motor and visual kernels and found that, across the population, the estimated motor kernel reflected inhibition in the early phase relative to swim times, and the estimated visual kernel reflected late-phase responses to visual motion feedback (**Fig. 3e, S2c**). Additionally, a post-swim ‘rebound’ appeared in the motor kernel (**Fig. 3e**) suggestive of a biophysical mechanism by which swimming first induces inhibition, followed by post-inhibitory rebound in serotonergic neurons. It also suggests that, during the rebound-like period, visual excitation causes the cells to fire proportional to visual speed. Although the biophysical mechanism of this increase in kernel values remains to be determined, we refer to this rise as the ‘rebound’. This suggests a mechanism by which the rebound by itself raises membrane potential, but not enough to cause spiking; similarly, visually-evoked excitation, alone, raises the membrane potential but not sufficient to trigger spiking; but when they coincide, they add and together cause voltage to rise above firing threshold, allowing serotonergic neurons to encode visual speed in a small time window following motor commands. Evidence of rebound in DRN neurons has been observed^32^. Rebound may occur through a variety of mechanisms^33–35^. Thus, a predictive model fit disentangles motor and visual contributions to the firing patterns of serotonergic neurons and suggests a mechanistic substrate for the way they transform motor-sensory inputs into spikes and eventually serotonin release.

We asked whether this temporal coincidence of the swim-evoked rebound and visually-evoked excitation is necessary for evoking spiking activity (**Fig. 3f, S2d**). Our previous work using calcium imaging indicated that DRN serotonergic neurons respond only to visual input in a precise window right after swims^14^ which, together with delays in visual input due to processing in visual centers, corresponds to the period when the fish receives visual motion feedback during late-phase swimming and the following coasting period (i.e., the short period when the body continues moving forward due to its momentum). This coincidence detection is represented as a Heaviside function in the LAE model which is 1 when its argument is positive and 0 otherwise (**Fig. 1f, 3f**), and may contribute to the fish’s ability to learn to attenuate swim vigor under high Gms condition only when visual motion is actively coupled to swimming, but not when the same motion patterns are passively replayed (**Fig. S3**). Thus we investigated the membrane potential and spiking activity of serotonergic neurons under three conditions: (1) swimming with visual feedback, (2) swimming without visual feedback, and (3) replayed visual feedback without swimming. Whereas swimming with feedback increased post-swim spikes, visual input without swimming elicited no change in spikes (**Fig. 3f, S2d**). Swimming without feedback caused the dip in firing rate during swimming, but did not elicitpost-swim spiking, while the membrane potential showed post-inhibitory rebound (**Fig. 3e, S2e**). Thus, through a mechanism of temporal coincidence detection, DRN serotonergic neurons effectively filter out sensory inputs that are irrelevant for motor vigor learning, and encode learning-relevant signals in post-swim spiking.

### The sources of inhibitory and excitatory inputs on the dendrites of serotonergic neurons

We sought to identify the pathways of motor- and visually-driven inputs^36^ that determine responses of raphe serotonergic neurons by using genetically-encoded neurotransmitter indicators (**Fig. 4**). Serotonergic raphe neurons receive both GABAergic and glutamatergic input^37^. Thus we expressed a GABA indicator (iGABASnFR)^38^ or a glutamate indicator (iGluSnFR)^39^ in *tph2*+ serotonergic neurons (**Fig. 4a,b**) and imaged their dendrites to monitor excitatory and inhibitory inputs during motor vigor learning. We observed a ‘wave’ of GABAergic input during swimming, whose amplitude increased when swim vigor increased at low Gms. This indicates that GABAergic input to serotonergic neurons during swimming encoded motor vigor during swimming (**Fig. 4c,d**). These GABAergic inputs likely originate from local GABAergic neurons in the DRN which encode the vigor of swimming **(Fig. S4a,b)**^14^ and may monosynaptically inhibit 5-HT neurons^40^. The timing of the GABA signal (**Fig. 4c**) was consistent with the timing of the decrease in the transmembrane voltage of serotonergic neurons (**Fig. 3d**). Conversely, glutamate input was present in two ‘waves’, the first wave occurring during swimming and the second one occurring immediately after swimming (**Fig. 4e,f** and **S4c,d**). The first wave reflected the motor command, or efference copy, and started slightly before swimming, and the second wave peaked after the swimming. The timing of the first glutamate wave coincided with the timing of the GABA signal suggesting that GABAergic inhibition masked the glutamatergic excitation in the early phase. The onset of the second glutamate wave coincided with the onset of the voltage ramp in serotonergic neurons (**Fig. 3d**), consistent with glutamate being the driver of the late-phase voltage increase. This second wave reflected the speed of visual feedback during swimming, which is proportional to Gms (**Fig. 4e,f**), consistent with the serotonin neurons’ spiking responses and serotonin release (**Fig. 1e**, **Fig 3d**). The timing of the glutamatergic signal was dependent on the delay of visual feedback (**Fig. S4e,f**). This second wave of glutamate, encoding visual input, remained intact even without swimming (**Fig. 4g,h**). This indicates that the observed dependence of visually-induced spiking on the presence of motor output (**Fig. 3f**) is due to mechanisms within the serotonergic neurons, and is not inherited from inputs upstream to the DRN.

**Figure 4.**
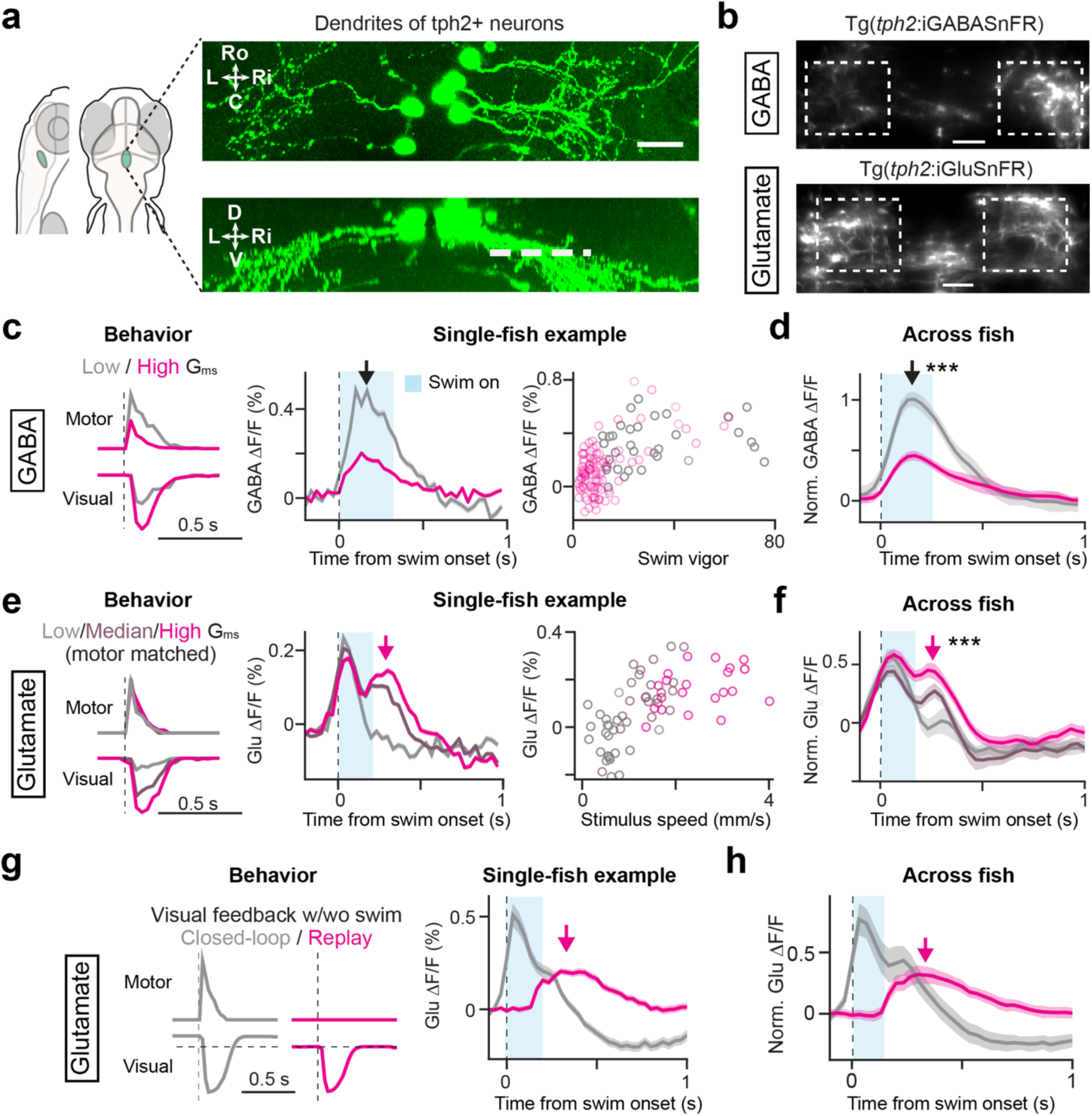
The sources of inhibitory and excitatory inputs on the dendrites of serotonergic neurons. **(a)** Morphology of raphe serotonergic neurons and their dendrites in zebrafish. Dorsal and side projections from a transgenic zebrafish that sparsely expresses GFP *Tg(tph2:Gal4; UAS:GFP)* are shown. The approximate axial location of dendritic imaging is indicated by a dashed line. Ro, rostral; C, caudal; L, left; Ri, right; D, dorsal; V, ventral. Scale bar, 20 µm. **(b)** Images of GABA indicator *Tg(tph2:iGABASnFR)* and glutamate indicator *Tg(tph2:iGluSnFR)* expressed in the dendrites of serotonergic neurons. Scale bar, 20 µm. **(c-d)** Dendritic GABA inputs during motor vigor learning. GABA signals encode swim vigor of individual swim bouts, with stronger GABA signals during strong bouts in low G_ms_. **c**, example from a single fish; **d**, summary of 12 sessions from 4 fish. Three different axial planes were imaged for each fish. Shades represent s.e.m. across experiments. ***, p < .001, paired signed-rank test for the peak amplitudes. **(e-f)** Dendritic glutamate inputs at various G_ms_. Data shown for swim bouts with various amounts of visual feedback randomly given at low, medium, or high G_ms_. We sub-selected swim events so that the average motor vigor equalizes among different G_ms_ conditions. The first peak of glutamate inputs encodes swim command (rising slightly before swim onset), and the second peak encodes the speed of visual feedback in the post-swim phase. **e**, example from a single fish; **f**, summary of 25 sessions from 9 fish. Two or three different axial planes were imaged for each fish. Shades represent s.e.m. across sessions. ***, p < .001, paired signed-rank test across all 3 paired G_ms_ groups for the second peak of the glutamate inputs. Data without the sub-selection of swim events are presented in Fig. S4c,d. **(g)** Visually-evoked glutamate inputs occur in the absence of motor actions. Example from a single fish. We sub-selected swim events so that the speed of visual feedback equalizes across conditions. Gray, closed-loop; magenta, open-loop replay trial with no swimming. Note that in replay, forward visual flow was replaced by zero visual flow to prevent swimming – see stimulus trace on the left – which is the likely cause for the more prolonged glutamate signal compared to the closed-loop stimulus. **(h)** Cross-fish average of glutamate inputs in response to closed-loop feedback during swimming (gray) or replay of visual feedback (magenta) shown in (g). N = 6 fish, 17 sessions. Shaded regions represent s.e.m. across sessions.

We next investigated the source of glutamatergic visual inputs into the raphe serotonergic neurons by identifying brain areas that respond to visual feedback during swimming and mapping their projection and innervation patterns in the brain (**Fig. S5**). Whole-brain neural activity imaging in swimming zebrafish under virtual reality setting with/without OMR-inducing stimuli and with/without visual feedback identified 6 major brain areas including the raphe that respond to visual feedback during swimming (**Fig. S5a-c**). Canonical correlation analysis between raphe serotonergic neurons and neural populations in other brain areas identified the thalamic area, which receives direct innervation from retinal ganglion cells^41^, as a potential input source (**Fig. S5d,e**). Consistent with this functional observation, we found that axonal projections from the thalamus area according to a single-cell projection atlas^42^ spatially overlap with the dendrites of raphe serotonergic neurons (**Fig. S5f**). Taken together, the above results of neurotransmitter imaging and identification of potential upstream brain areas suggest that neural activity of raphe serotonergic neurons is determined by swim-evoked inhibition from local GABAergic neurons and visually-evoked glutamatergic excitation from remote upstream areas.

To test whether such a model explains the data, we formulated an artificial neural network that simulates GABAergic neurons and serotonergic neurons in the raphe nucleus that receive swim- and visually-evoked inputs, respectively (**Fig. 5a,b**). We also considered other network architectures that could explain behavior, but closer inspection showed them to be inconsistent with the spiking and glutamate/GABA signals (**Fig. S6**). GABAergic neurons inhibit serotonergic neurons^40^. Based on the estimated response kernel derived from voltage imaging data (**Fig. 3e**) and membrane potential dynamics following swim bouts in the absence of visual input (**Fig. S2d,e**), we included hyperpolarization-induced cation channels in the serotonergic neurons. Simulating this system as a spiking neural network driven by swimming and visual motion (**Fig. 5c**) generates a timeseries of membrane potential and spiking activities (**Fig. 5d,e**) that can be compared to the experimental data.

**Figure 5.**
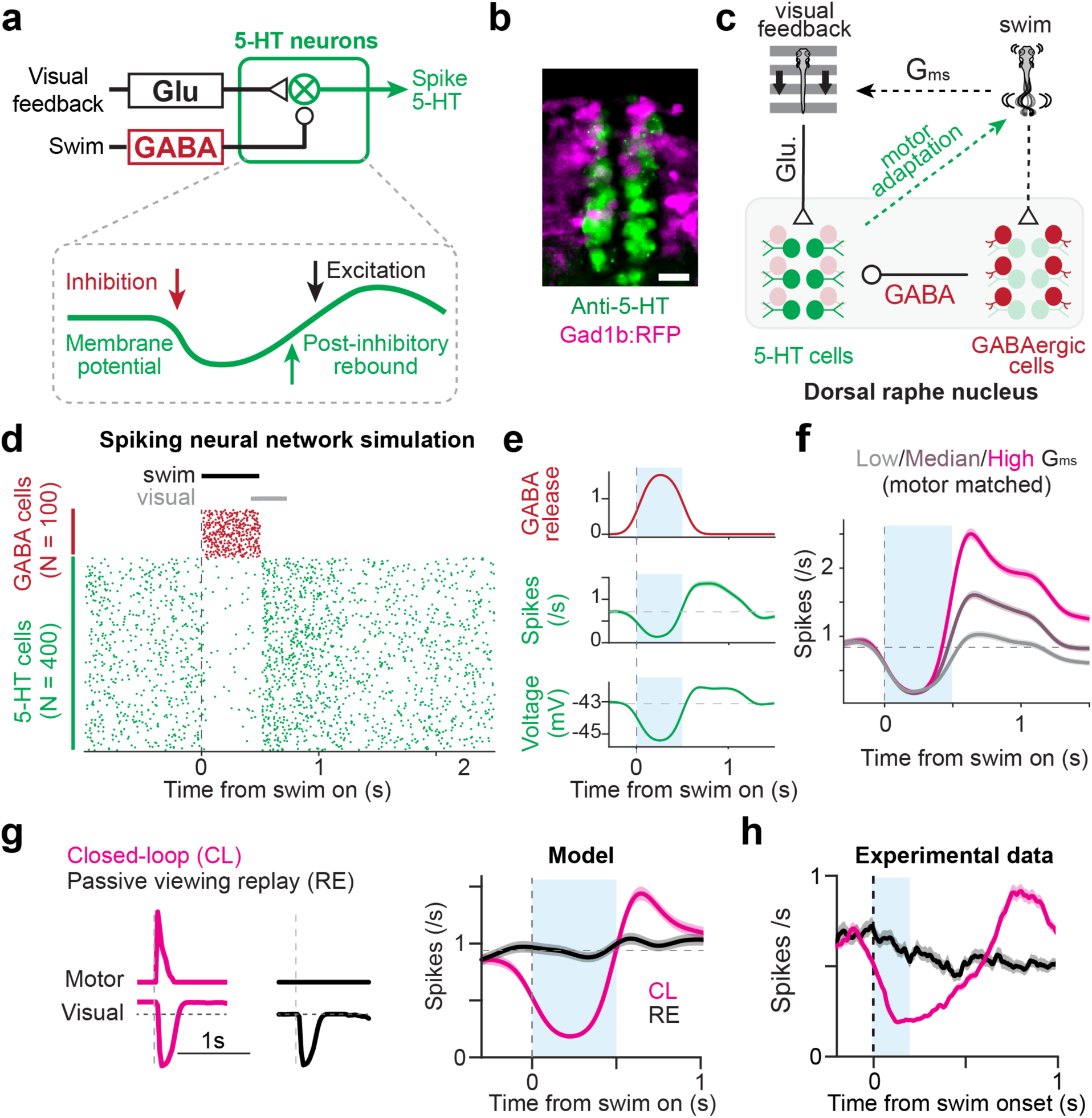
Simulation of sensory-motor integration in the serotonergic raphe using spiking neural network. **(a)** Model schematic showing GABAergic motor-command-encoding input, glutamatergic visual input, post-inhibitory rebound, and spike/5-HT output. **(b)** Serotonergic (green) and GABAergic neurons (magenta) in the raphe nucleus, which are largely non-overlapping populations in larval zebrafish. Serotonergic neurons were visualized using immunostaining for 5-HT in a transgenic zebrafish that express RFP in GABAergic neurons (see Methods). Scale bar, 10 µm. **(c)** Structure of spiking neural network model. **(d)** Simulated spiking activity during a single swim bout. **(e)** Resulting spiking, membrane potential, and GABA activity signals in the model during a swim bout. **(f)** The model shows during-swim suppression and post-swim visual response in serotonergic neurons, which recapitulates experimental results (see Fig. 3d). **(g)** The simulated serotonergic neurons do not show visual response without motor-induced GABAergic inhibition. **(h)** Experimental data, without swim bout, does not show visual response after swimming, qualitatively matching the model. Replot of the same data in Fig. 3f.

This model reproduced the spiking patterns and the encoding of the velocity of swim-induced visual feedback seen in experimental data, and post-swim spike rates increased proportionally to Gms (**Fig. 5f**). We also examined whether the model can reproduce the lack of spiking response in serotonergic neurons to replayed visual motion uncoupled to swimming by removing GABAergic neurons from the model. Due to the loss of post-inhibitory rebound, spiking responses to visual feedback was obliterated in serotonergic neurons, which qualitatively matches the absence of visual responses when the fish is not swimming in experimental data (**Fig. 5g,h**).

Taken together, the spiking network model and its qualitative match to experimental data suggests that synergetic effects of swimming-evoked, post-inhibitory rebound and visually-evoked glutamatergic input form the mechanistic implementation of robust encoding of action effectiveness while filtering out irrelevant sensory inputs in DRN serotonergic neurons.

### GABAergic inhibition enables the encoding of action effectiveness in serotonergic neurons

Serotonergic neurons in the DRN receive inhibitory input form local GABAergic neurons^43,44^. In zebrafish, raphe GABAergic neurons encode the vigor of swimming (**Fig. S4a,b**)^14^ and likely contribute to the swim-evoked inhibition of serotonergic neurons that we observed using voltage imaging (**Fig. 3**). The ensuing post-inhibitory rebound may provide a time window for visual input to evoke spiking in serotonergic neurons (**Fig. 6a**).

**Figure 6.**
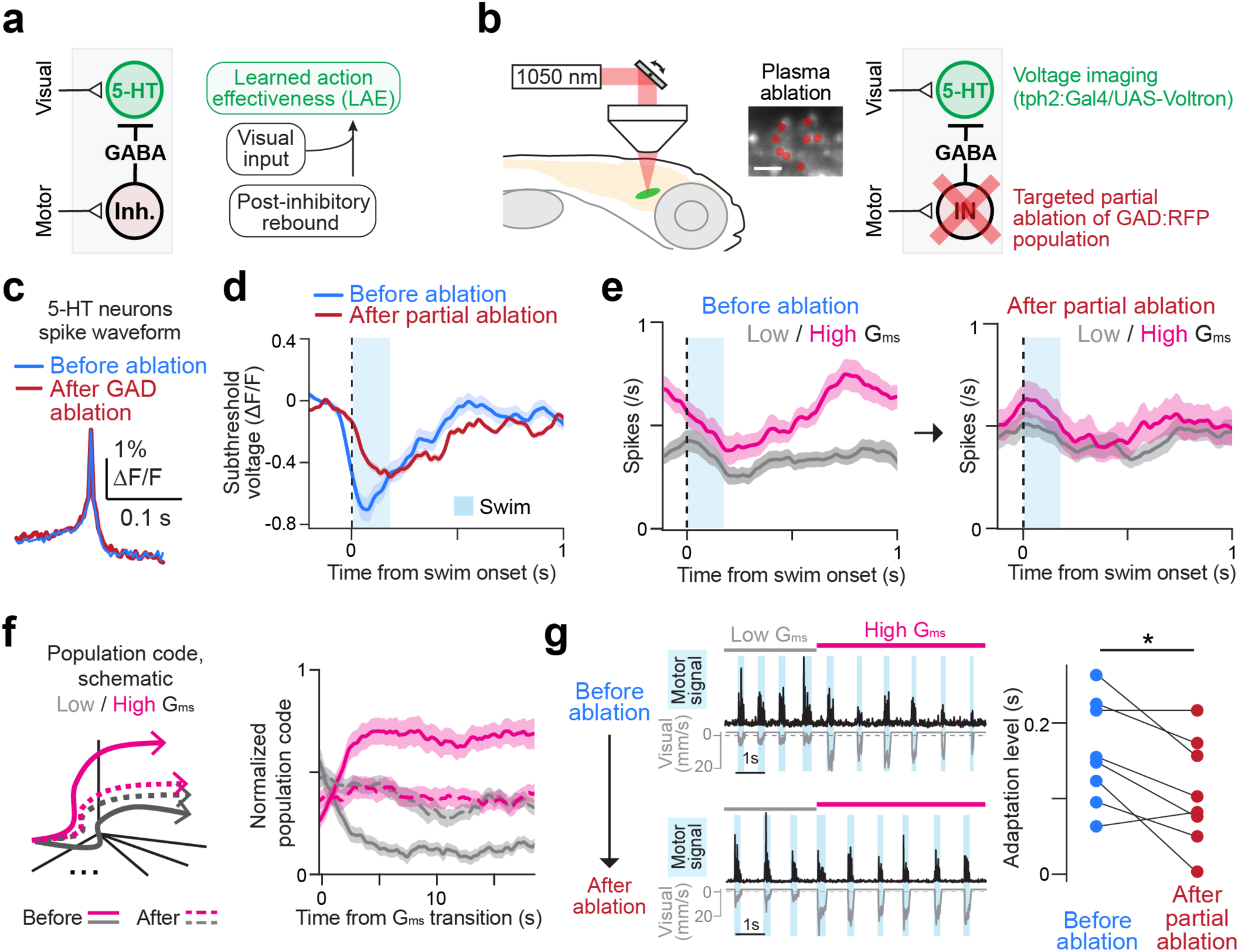
GABAergic neuron ablation impairs DRN motor-gating and population coding. **(a)** Hypothetical circuit diagram in the raphe nucleus. Learned action effectiveness (LAE) is encoded by a synergistic action of swim-induced post-inhibitory rebound and visual inputs. **(b)** Ablating a subset of GABAergic neurons inside the raphe (ablation) and outside the raphe (control ablation) using a two-photon plasma ablation method. We used the wavelength of 1050 nm to ablate GABAergic neurons that express RFP (center). The loci of the ablation and control ablation are shown in Fig. S7a,b. We performed voltage imaging using Voltron in tph2+ neurons before and after the ablation. Scale bar, 10 µm. **(c)** Spike waveform in an example serotonergic neuron remains intact after the ablation, providing evidence that those neurons are unharmed by the ablation of nearby GABAergic neurons. **(d)** Changes of membrane potential dynamics in serotonergic neurons before and after the partial ablation of the GABAergic population. The ablation of raphe GABAergic neurons decreases swimming-evoked inhibition, consistent with the model. It also decreases post-swim excitation, also consistent with the post-inhibitory rebound component of the model. N = 9 fish; 48 high G_ms_ neurons; shaded area represents s.e.m. across cells. **(e)** The partial ablation of the raphe GABAergic population results in the loss of visual feedback encoding. *Left*, spiking in serotonergic neurons before ablation during low (gray) and high (magenta) G_ms_. *Right*, spiking in 5-HT neurons in the same animals after the ablation. Activity no longer encodes the speed of visual feedback depending on G_ms_, while the baseline firing remained intact. Data from 48 high G_ms_ neurons in 9 fish; shaded area, s.e.m. across cells. **(f)** The ablation of DRN GABAergic neuron leads to failed population coding of LAE in serotonergic neurons. *Left*, schematic of population coding for G_ms_. A version of linear discriminant analysis based on spiking activity is performed to maximally separate trajectories between low vs. high G_ms_ (see **Methods**). *Right*, in intact fish, the 5-HT neuronal population code reliably distinguishes low from high G_ms_ (solid gray and magenta lines). After ablation, the population code no longer reliably distinguishes between low vs. high G_ms_ (dashed lines). N = 8 fish, 15 sessions, 10-20 cells simultaneously recorded per session; shaded area, s.e.m. across sessions. **(g)** The ablation of DRN GABAergic neurons leads to a reduction in motor adaptation. Adaptation level is the time within swim bouts during which a statistical difference exists between swim signal in low vs. high G_ms_ (see **Methods**). N = 8 fish. *, p = .0277, paired signed-rank test.

We therefore investigated whether targeted ablation of raphe GABAergic neurons impairs the encoding of visual feedback during swimming in serotonergic neurons and affects motor adaptation behavior (**Fig. 6**). We used two-photon plasma ablation^45^ of raphe GABAergic neurons that express red fluorescence proteins (**Fig. 6b**) and performed voltage imaging in serotonergic neurons before and after the ablation to examine how the ablation affects their membrane potential modulation and spiking activity during motor adaptation. The ablation of raphe GABAergic neurons did not alter the spike shapes of serotonergic neurons, indicating minimal collateral damage (**Fig. 6c**). As a control, we also performed ablations of GABAergic neurons outside the raphe (**Fig. S7a,b**). We found that the ablation of a subset of raphe GABAergic neurons led to a reduction in inhibition of serotonergic neurons during swimming (**Fig. 6d**), which was not observed by the control ablation (**Fig. S7c**), indicating that the raphe GABAergic neurons contribute to swim-evoked inhibition of serotonergic neurons.

Specifically, our simulation (**Fig. 5**) predicts that the loss of swim-evoked inhibition should lead to a loss of spiking response to visual feedback during swimming. We found that the visually-evoked increase in spike rate in the post-swim period was reduced after the ablation (**Fig. 6e**), leading to a complete loss of encoding of learned action effectiveness at the population level (**Fig. 6f**; Methods). Such changes were only partially observed in the control ablation (**Fig. S7d,e**). We also observed significant impairment of motor vigor learning after the ablation of raphe GABAergic neurons (**Fig. 6g**), which was not observed after the control ablation (**Fig. S7f**). Taken together, these results indicate that GABAergic inhibition of serotonergic neurons is core to both the learning of action effectiveness in the serotonergic system and that such neural circuit motifs in zebrafish may have broader implications in how the serotonergic system integrates multiple streams of information to control behavior in vertebrate species.

## Discussion

The precise multimodal measurements of dendritic inputs, somatic voltage dynamics and neuromodulatory outputs reported here provide a rare view into the inner circuit and biophysical workings of the vertebrate serotonergic raphe during learning behavior. The serotonergic system has been implicated in many neural processes including sensory responsiveness^46,47^, sleep regulation^48^, motor learning and planning^12,13,49–51^, mood regulation^18,52,53^, and shifts between interoception and exteroception^54,55^. Despite the known involvement of the serotonergic system in this wide range of processes, the inner workings of the serotonergic system^56,57^ have remained underexplored at the level of cellular and circuit mechanisms of neuronal dynamics and computation.

Building an understanding of neural computation involves identifying the input-output properties of the involved neurons. This task can become complex when a computation involves multiple brain regions and cell types, where it can be challenging to disentangle response properties that are inherited from other regions from those that result from information processing within the region^58^. This question can be especially hard to address in nuclei such as the serotonergic raphe, in part because it is hard to access its large and cell-type specific populations in a small nucleus deep beneath the cortex, where most of the recordings in mammals were done through electrophysiology or fiber photometry^53,59,60^. In light of the development of modern imaging sensors^61,62^ and other whole-brain imaging technology^63–65^ one can record multiple types of neuronal signals in the zebrafish raphe that are normally inaccessible. Our study provides a systematic way to track upstream inputs, local computation, and downstream outputs of the serotonergic system using multimodal imaging. These signals show dynamics inside the DRN that underlie the computation of action outcomes and suggest that the global effects of neuromodulation arise from a precise, fast-timescale, cellular and circuit mechanisms, and that the nuclei themselves implement these computations rather than simply inherit and relay them from upstream input. Future multi-color variants of the molecular and voltage indicators will make it possible to image multiple signal modalities simultaneously.

Motor learning relies on the animal’s ability to identify the causal outcomes resulting from ones actions amidst numerous streams of ongoing sensory input, and temporal coincidence detection can facilitate this process. Coincidence-detection mechanisms have been reported across various domains within neuroscience, such as in dendritic integration^66^, synaptic plasticity^67^, and sensory processing^68^. Synergetic effects of post-inhibitory rebound with excitatory inputs, whereby rebound brings a neuron closer to firing threshold so that excitatory inputs can drive it to spiking more easily, have been proposed as a coincidence detection mechanism in auditory processing^26,27^ and associative learning^69^. Our study, utilizing both empirical data and a spiking network model, provides evidence that post-inhibitory rebound may mediate robust encoding of action effectiveness for motor learning. This mechanism effectively aligns action causes with sensory outcomes and appropriately discounts sensory feedback that was less likely to be related to the actions (**Fig. 5**). Testing the rebound hypothesis beyond the model-based results presented here will require further experiments such as whole-cell intracellular recording^70^ and transcriptomics to identify the responsible membrane ion channels, which forms a promising future direction of research.

The serotonergic system controls various brain states^18,19,48–50,59,71–76^, yet its internal computational principles require clarification. We propose that the computational motif that enables action-outcome computation in the serotonergic system in zebrafish may generalize to computations that trigger other brain state transitions. For example, action effectiveness can reflect levels of challenge in a variety of situations including in learned-helplessness contexts^13,76,77^. Our study indicates that serotonin levels decrease as a function of the difficulty of the task; the input-output transform within the serotonergic system may thus reflect an internal ‘credit system’ for the outcomes of actions, which may be related to reward encoding in the serotonergic system^15,78^, or more generally, the encoding of the current overall level of challenge or perseverance in the environment in broader behavioral contexts^19,75^. *Action effectiveness* within the zebrafish motor learning model considered here may relate, conceptually, to *self-efficacy* as used in human psychology, which is defined as one’s belief in one’s capacity to act in the ways necessary to reach specific goals^79,80^.

We have formulated a motor learning framework based on the concept of learned action effectiveness (LAE). This computational framework effectively accounts for motor learning at the behavioral level, serotonin release dynamics, and population coding in the serotonergic system during short-term motor learning. Additionally, we have constructed a spiking neural network implementation for LAE, which facilitates a form of sensory-motor credit assignment in the form of coincidence detection, mediated by local GABAergic neurons. This neural network model allows for insights into neurophysiological data, including spikes, membrane potential, and neurotransmitter dynamics, at millisecond timescales. Our findings highlight that serotonergic neuromodulation relies on fast, precise circuit computations within the raphe, such as action-outcome coincidence detection, to directly influence behavioral adaptation. This suggests that similarly intricate mechanisms may operate across other neuromodulatory nuclei. These mechanisms, though likely to vary in specific details, may serve as a general framework for neuromodulation, enabling different systems to integrate diverse inputs with temporal precision to modulate behavior. Further exploration of cellular and circuit dynamics within these nuclei during behavior in multiple species, using genetically-encoded sensors for voltage, neurotransmitters, and neuromodulators will deepen our understanding of the common principles underlying the dynamic regulation of brain states by different neuromodulatory systems.

## Acknowledgements

We thank Carsen Stringer, Yu Mu, Florian Engert, Sandro Romani, Brett Mensh, and David Prober for their comments on the manuscript. We thank Liam Paninski for discussions on data analysis. We thank Adam Cohen, Brett Mensh, and members from Ahrens lab and Kawashima lab for discussions on this project. This project was funded by the Howard Hughes Medical Institute (M.B.A.), the Simons Foundation Simons Collaboration on the Global Brain Award 542943SPI (M.B.A.), the Israel Science Foundation Individual Grant 688/22 (T.K.), the Binational Science Foundation #2021746 (T.K.), an Azrieli faculty fellowship (T.K.), an Abisch Frenkel Foundation (T.K.), Jonathan and Joan Birnbach (T.K.), Dan Lebas & Ruth Sonnewend (T.K.), an Internal grant from the Center for New Scientists at Weizmann Institute of Science (T.K.), the Israeli Council for Higher Education (CHE) via the Weizmann Data Science Research Center (R.H.), the Moshe Meir Horwitz Fund (R.H.), Max Planck Society (H.B.), Bridge Position Program for Advancing Women in Science and Gender Equality in Weizmann Institute of Science (I.S.), and Alexander von Humboldt postdoctoral fellowship (I.S.).

## Author contributions

TK conceived the project, performed experiments on motor adaptation behavior, voltage imaging, neurotransmitter imaging and two-photon plasma ablation, performed data analysis, and wrote the manuscript. ZW conceived the project, developed data processing pipelines for voltage imaging and neurotransmitter imaging, performed data analysis and modeling, and wrote the manuscript. RH performed the serotonin imaging experiment. SN generated transgenic zebrafish for voltage and neurotransmitter imaging. IS and HB generated transgenic zebrafish for serotonin imaging. MBA conceived and supervised this study, analyzed data, and wrote the manuscript.

## Methods

### Zebrafish husbandry

All zebrafish experiments were approved by the Institutional Animal Care and Use Committee (IACUC) at Janelia Research Campus, HHMI and the Institutional Animal Care and Use Committee (IACUC) and the Institutional Biosafety Committee (IBC) of the Weizmann Institute of Science and by the Israeli National Law for the Protection of Animals - Experiments with Animals (1994). Larvae were kept at a 10 h dark, 14 h light cycle at 28°C. Larval sex is not specified at this developmental stage and was therefore not determined.

### Transgenic zebrafish

Transgenic zebrafish that pan-neuronally express genetically-encoded serotonin indicator *Tg(HuC:iSeroSnFR)* (**Fig. 1**) was generated in AB strain with Casper mutation^81^ as described elsewhere^29^. To enhance the brightness, we use homozygous, in-crossed larvae at the age of 5-6 dpf. Transgenic zebrafish that express genetically-encoded voltage indicator, Voltron, (**Fig. 2, 3**) were obtained by crossing transgenic zebrafish that express Voltron under UAS promoter *Tg(UAS:Voltron)*^jf42Tg^ ^30^ with transgenic zebrafish that express Gal4 transactivator under the tph2 promoter *Tg(tph2:Gal4)*^y228Tg^ ^47^. To label Voltron-expressing neurons with the accompanying fluorescent dye, 3-day old embryos were incubated in dye solution [3.3 μM JF_525_-HaloTag ligand^82^ and 0.3 % DMSO] in fish rearing water at room temperature for two hours. After screening for the fluorescence of the JF dye in the brain, the fish were returned to fish rearing water with food until the time of the experiment.

Transgenic zebrafish expressing a GABA indicator, iGABASnFR^38^, or a glutamate indicator, SF-iGluSnFR^39^, (**Fig. 4**) were generated by cloning these genes in the direct downstream of the tph2 promoter^47^ on a Tol2 vector. These plasmids were injected into two-cell stage embryos of AB Casper mutant zebrafish with mRNA of Tol2 transposase to generate founder transgenic zebrafish. Experiments were performed using 5-6 dpf embryos generated by out-crossing F0 founder to AB Casper fish.

GABAergic neurons in the DRN (**Fig. 5b**, **Fig. 6**) were labelled using a transgenic zebrafish that expresses RFP in *gad1b*+ neurons *Tg(gad1b:loxP-RFP-loxP-GFP)*^nns26Tg^. The anatomical co-labeling with serotonergic neurons (**Fig. 5b**) used an anti-5-HT rabbit polyclonal antibody (S5545, Sigma-Aldrich) for immunostaining as described in our previous work^14^.

### Preparation for zebrafish imaging experiments

Imaging experiments were performed using 5- or 6-day larval zebrafish. The zebrafish was immobilized and mounted to an imaging chamber as described previously^14^ with minor modifications. To detect small fluorescence changes of voltage and neurotransmitter indicators in the soma and dendrites of DRN serotonergic neurons (**Figs. 2, 3, 4, 6, S2, S4, S7**), we needed to stop the circulation to avoid shadowing effects of blood cells that pass through the excitation beam during the imaging^30^. For this purpose, the zebrafish were habituated in an artificial cerebrospinal fluid (ACSF) [in mM: 120 NaCl, 2.9 KCl, 2.1 CaCl2, 1.2 MgCl_2_, 20 NaHCO_3_, 1.25 NaHPO_4_, 10 Glucose] pre-bubbled with carbogen gas (95% O_2_, 5% CO_2_) for 30 minutes. The muscle of the zebrafish was then paralyzed by a short (up to 30 seconds) bath incubation with alpha-bungarotoxin (1 mg/ml, Thermo Fischer Scientific, B1601) dissolved in an external solution. After the fish became immobile, heart movement of the zebrafish was stopped by microforceps to disrupt circulation. The zebrafish showed robust optomotor behavior in pre-bubbled ACSF for several hours. The zebrafish was further mounted to a custom-made chamber using 2% agarose (Sigma-Aldrich, A9414) and placed under a light-sheet microscope^65^ with a 20x objective lens (Olympus, XLUMPLFLN). For serotonin imaging (**Fig. 1**), we performed experiments in normal fish rearing water without using ACSF and heart surgery because of the lack of blood shadow artifact in the dorsal hindbrain.

### Light-sheet imaging

Imaging was performed in a light-sheet microscope according to a published design^65^ with modifications targeted at optimizing Voltron imaging. To increase the fraction of time during which the imaged cells were illuminated by the excitation laser beam, the beam was expanded in the horizontal dimension using a pair of cylindrical lens (LJ1878L1-A (f=10mm) and LJ1402L1-A (f=40mm), Thorlabs). Imaging was performed using a 488 nm excitation laser (80 μW), a 562/40 emission filter for Voltron (Semrock, FF01-562/40) or a 525/50 emission filter for neurotransmitter indicators (Semrock, FF01-525/50), and a sCMOS camera (Hamamatsu, ORCA Flash4.0 v2). We used a frame rate of 300 frames/second for voltage imaging, and 30 frames/second for neurotransmitter indicators. In this setup, the pixel dimension on the camera was 0.293 μm/pixel and the imaged neurons occupied an area of 150-200 pixels on the image.

### Behavioral Assays in Virtual Reality

We simulated an environment in which the animals swim along a one-dimensional virtual track consisting of red and black bars (each 2 mm thick) perpendicular to the direction of swimming (**Fig. 1b**) as previously described^14,21,65^. The visual effect of swimming was mimicked by accelerating the gratings backward when a fictive swim bout was detected. The visual environment slowly moved forward at the speed of 2 mm/s between swim bouts to visually simulate backward movement of the fish due to the virtual backward water current. The fish swim against this virtual water current to stabilize their positions^23^. Thus the forward velocity of the gratings is programmed as:

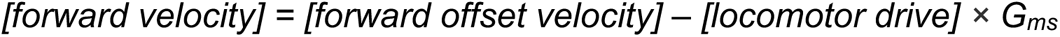

(**Fig. 1b**). During strong swim bouts, the visual environment moves backward simulating forward swimming. G_ms_ is a parameter controlling the amount of backward motion arising from fictive swimming and simulates action effectiveness.

We recorded swim signals (fictive swims) from the axonal bundles of spinal motoneurons in the tail by attaching a pair of large barrel electrodes to the left and right side of the dorsal part of the tail. Signals were amplified by an amplifier (Molecular Devices, AxoClamp 700B or a combination of Intan RHD2132 chip and RHD2000 evaluation board) and recorded at 6 kHz using custom software written in C# (Microsoft) or in Python. For synchronization between the swimming signals and neural activity imaging, camera trigger signals that initiate the acquisition of individual frames in the light-sheet microscope were recorded simultaneously with the swim signals. Signals from the electrodes were processed and individual swim events were detected according to a method described previously^14^. Briefly, the raw signals were high-pass filtered, squared and smoothed by applying a Gaussian filter (σ = 3.3 milliseconds). The resulting traces were defined to be the fictive swim signal (**Fig. 1b**). Individual swim bouts were detected by finding the time points at which the swim signal crossed a threshold. This threshold was automatically set to lie just above a noise level based on a histogram of the swim signals^21,65^. Visual stimuli were delivered using a DLP projector (Picopix PPX-4010, Philips or LV130, Optoma) which have low input lag of 16 milliseconds. The delay from fictive swim acquisition to the display of the gratings with the programmed velocity is about 35 ms in our virtual reality setup.

### Behavioral tasks

#### Motor vigor learning

We examined the fish’s behavior, neural responses, and neurotransmitter release during the motor vigor learning task (**Fig. 1, 2, 3a-e, 4c, 5, 6, S1a,b, S2b, S4c,d, S7c-f**), where fish receive visual feedback from swimming in closed loop and adapt their swim vigor to the motosensory gain G_ms_. We alternated G_ms_ between low and high levels, each lasting 20 seconds. The value of high G_ms_ was chosen to be two to three times higher than that of low G_ms_ ^21^, and the value of low G_ms_ was set to the lowest value at which the zebrafish can stabilize their position in the virtual reality arena.

#### Visuomotor decoupling

We measured the neural activity of serotonergic neurons and glutamate release on their dendrites when the fish’s swimming and visual velocity input resembling feedback are temporally decoupled (**Fig. 3f**, **Fig. 4g,h, Fig. S2d,e**). We designed three types of visuomotor events: closed-loop swimming (CL), swimming without any feedback (open-loop, OL) and replay of visual feedback without swimming (RE). In CL events, visual feedback was provided when the fish swam, using high G_ms_. In the OL event, G_ms_ was set to zero which resulted in no visual feedback in response to swimming. In the RE event, we replayed a backward visual motion pattern recorded in the preceding closed-loop epoch and only analyzed events that did not overlap with swimming. The latter two types of decoupling events were randomly presented scattered among the closed-loop events. OL events occasionally triggered struggling-like swimming^83^. We excluded such events from the analysis and subsampled events with similar swim vigor as those in CL events. For RE events, we subsampled those that did not coincide with swimming.

We tested each fish in the motor adaptation task for 5 minutes before testing them in this task to ensure its capacity for responding to visual feedback and adapting their swim vigor.

#### Random gain/delay

We examined glutamate release on the dendrites of serotonergic neurons in the conditions where we [i] changed G_ms_ randomly at three levels (low, medium, high) for individual swim bouts (**Fig. 4e,f**) or [ii] delayed the visual feedback by three lags (0 ms, 200 ms, 400 ms) (**Fig. S4e,f**). In the former task of varying G_ms_, the medium G_ms_ was set to the average value of low and high G_ms_. In the latter task of varying delays, the delay was introduced randomly in 10% of the swim events for each of the delayed conditions. We tested each fish in the motor adaptation task for 5 minutes before testing it in this task to ensure its capacity for responding to visual feedback and adapt its swim vigor.

#### Short-term motor learning

This behavioral assay for testing the persistent learning effect (**Fig. S1d-f**) was previously described^14^. Briefly, the paradigm consisted of a 15-second initialization period with low G_ms_, a 7-second or 30-second training period with high G_ms_, a 10-second delay period with stopped visual stimuli, and a 5-second test period with medium G_ms_. We performed 20 - 30 trials per recording session. The data in Fig. S1d-f was acquired in the above study and re-analyzed for this study to compare with the prediction of LAE model.

We also tested a variant of this behavioral assay to de-couple swimming and visual feedback (**Fig. S3**) to investigate whether training with visual feedback alone without swimming for 30 seconds have similar learning effect to the above motor learning task. In half of the trials, visual feedback during the preceding 30-second training with high G_ms_ was replayed during the training period. To prevent the occurrence of swimming during the replay, only the backward component of visual stimuli motion was replayed.

### Processing pipeline for voltage imaging data

We built a data processing pipeline for voltage imaging data to automatically perform camera-noise correction, motion correction, denoising, and cell segmentation (**Fig. 2b**).

#### Step #1: Camera-noise correction

We adopted the camera noise correction algorithm from refs. ^84,85^. The pixel readout is computed as

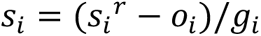

where *s*_i_^*r*^ is the raw camera readout at pixel i and *s*_i_ is the corrected one; *o*_i_ is the camera offset and *g*_i_ is the camera gain at pixel i. We estimated the offset *o*_i_ and baseline variance *v*_i_in the no light condition as the mean and variance of 60k images. The gain for each pixel was calculated as

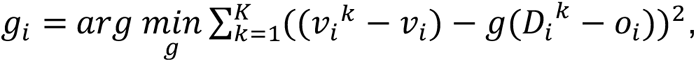

where K is the total number of illumination levels acquired, k is the k^th^ illumination level, *D*_i_^*k*^and *v*_i_^*k*^are the mean and variance of the 20k images acquired in the k^th^ illumination level. In our setup, we measured them by varying the laser power from 0mW (no light condition) to 18 mW. Empirically we found that the camera-noise correction can improve motion correction and other later steps in the processing^85,86^.

#### Step #2: Motion correction

We performed the two-dimensional rigid registration of the images using a custom Python script based on the dipy package^87^, to correct drifts in the sequentially recorded images at the subpixel level.

#### Step #3: Image-series denoising

We performed image-series denoising for motion-corrected video Y by finding its low rank (at K) representation

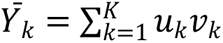

with residual *R*_*k*_ = *Y* − *Ȳ*_*k*_ is statistically white (within 99% confidence interval). Here, *u*_*k*_, *v*_*k*_ are spatial and temporal components respectively and determined by the objective

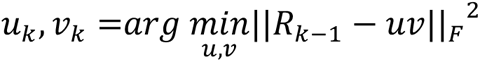

where F denotes Frobenius norm. This is equivalent to performing iterative principal components analysis (PCA) on Y and stopping the iteration at the k^th^ components as the residual is closed to the white noise. Since the number of pixels is large which creates difficulty in computing PCA, we alternatively divided whole images into overlapped small patches (namely local PCA), performed this denoising procedure for each patch, and stitched the denoised patches back to a full denoised movie. Moreover, since video Y can be presented as a 3D tensor, we also tested the tensor-based optimization as

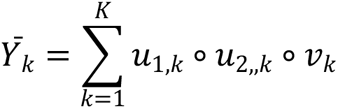

where *u*_(,*k*_ and *u*_2,,*k*_ are two spatial components along the vertical and horizontal directions of the imaging, respectively. We found that tensor decomposition would generate stripe-like spatial correlation horizontally (which might stem from the power fluctuation of the horizontal light-sheet illumination), and thus used local PCA instead throughout the denoising step.

#### Step #4: Cell segmentation

We performed cell segmentation on the denoised movie (*Ȳ*_*k*_) using semi-nonnegative factorization, where

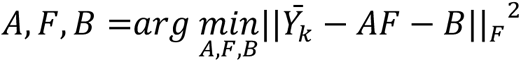

such that

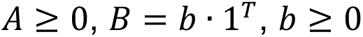

where A is the matrix of the nonnegative components in which the pixels were connected in space (ROIs); F is the matrix of the temporal components and presents the fluorescent dynamics of ROIs, and B is the temporally-constant background. We initialized the components using super-pixels^88^, which include the local pixels with neighboring correlations larger than 0.8. We then computed

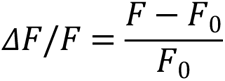

for each component (or ROI) where *F*_0_is baseline fluorescence computed as a running 20% percentile of F within a 3-minute time window. In our hands, denoising step does not affect dynamics of spike generation (thus spike detection) or membrane potential fluctuation, comparing to the compute of ΔF/F using hand-draw ROIs. Since Voltron is a negative response indicator, we used the flipped signal (−ΔF/F) for subsequent analyses for spike detection and subthreshold activity estimation.

#### Step #5: Spike detection

We trained a machine-learning neural network (**Fig. 2c**) in order to automatically extract the spikes from the time series data of ΔF/F. The network consists of two LSTM layers and a dropout layer which interconnects them. To train this neural network, we used simultaneous electrophysiology and imaging of cerebellar neurons expressing Voltron acquired in our previous work^30^. The goal of the network is to determine whether there is a spike event in the middle of the time series (the length of the time series is 41 frames or 136.67 ms) by minimizing the loss function of the cross-entropy between the prediction of spike from voltage imaging with the ground truth from the electrophysiology (**Fig 2c**; gray dots, ground truth from electrophysiology; black, prediction of the network). We performed the 10-fold cross-validation in network training. We applied the trained network model to extract the spikes automatically thereafter.

#### Step #6: Subthreshold activity estimation

We estimated the subthreshold activity as a rolling median filter of time series data of ΔF/F with window size at 70 ms. To avoid the nonlinearity of spikes (a depolarization followed by a repolarization), we clipped out the frames (from -1 to +1) around detected spikes while running the median filter.

### Processing of neurotransmitter imaging data

We adapted parts of the voltage imaging processing pipeline to automate the neurotransmitter and neuromodulator imaging processing, which included camera noise correction, then motion correction, and denoising (**Fig. 2b**).

#### Identification of the regions of interests and estimation of dynamics

After we obtained the denoised movie (which followed the same procedure as that in Voltron imaging processing, **Fig. 2b**), we identified the super-pixels^88^, which includes the local pixels with neighboring correlations larger than 0.8. We computed average fluorescence F inside all super-pixels (weighted equally) and then computed

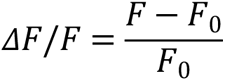

for each movie, where *F*_0_ is baseline fluorescence computed as a running 20% percentile of F within a 3-minute time window.

### Statistical analysis

All statistical analyses were performed in python using the Scipy package^89^. Statistical details of experiments, including the types of statistical tests and sample numbers can be found in the figure legends. Briefly, we applied nonparametric statistics for comparisons: a signed-rank test for paired comparison and a rank-sum test for unpaired comparison, and Kruskal–Wallis test for unpaired comparison across more than 2 conditions. We used Spearman correlation to measure the correlation of neural variables with behaviors.

### Behavioral quantifications

#### Inclusion criteria

Zebrafish that do not show robust adaptation of swim vigor to the change of G_ms_ may have problems in perceiving the visual stimuli projected below the fish (**Fig. 1b**) or developmentally defective. Therefore, we excluded fish whose swim vigor during high G_ms_ was not statistically lower than those during low G_ms_ in the motor vigor learning, with a cut-off threshold of p<0.01 using one-side rank-sum test.

#### Equalizing motor outputs at different G_ms_

To disentangle the neural encoding of swim vigor and visual feedback, we subsampled the swim bouts with similar vigors at different G_ms_, to analyze *“motor-equalized trials”* (**Fig. 3d, 4e**) as follows. We first computed the empirical cumulative distribution (*ECD*) of swim vigors across all swim bouts at a given G_ms_. We then decided the overlapped range of swim vigors to subsample according to *ECD*s. Finally, we tweaked the subsample range for each G_ms_ to make sure that the average swim vigor of the subsamples were statistically identical across G_ms_ (Kruskal–Wallis test, p<0.05). We applied this procedure to subsample trials in tasks of motor adaptation (**Fig. 3d**), random G_ms_ (**Fig. 4e**) and random visual-feedback delays (**Fig. S4e**).

#### Adaptability of swim patterns

We quantified the ability to perform motor vigor learning before and after ablation of DRN GABAergic neurons or control GABAergic neurons (**Fig. 6g, S7f**) as follows. We binned the swim vigor data at 300 Hz, and generated a vector of time series of swim vigor for 1 second, which is long enough to cover most swim patterns. We filled post-swim points in the vector as zeros. We then compared the moment-by-moment difference of swim vigor between low and high G_ms_. The cumulative period of time, where swim vigor was larger (p<0.05, one-tailed rank-sum test) at low G_ms_, was used to calculate the “adaptation level” of swimming between G_ms_.

### Kernel fits of membrane potential and spikes to behavioral variables

Response kernel estimation of serotonergic neurons based on behavioral variables (**Fig. 4e**) was performed as follows. We predicted spike events and subthreshold membrane potentials from the behavioral variables of time series of the swim vigor

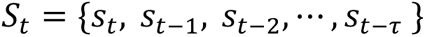

and that of the visual input

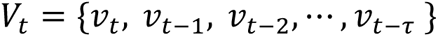

and that of the recent spike events

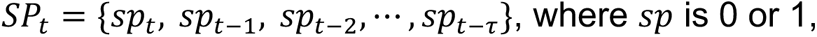

at a given time *t*, allowing history dependence (*t-τ*) up to 300 time points (1 second). Since spiking activity is sampled at 300 Hz, for each single frame, the spike event is binary, we thus modeled the spike event as

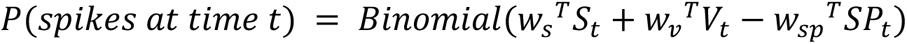

We optimized this probabilistic model from the data using generalized linear model with logit link function and a *l*_2_-form regularization on kernels of the swim vigor, i.e. *w*_*s*_, the visual input, i.e. *w*_*v*_, and the spike history, i.e. *w*_*sp*_. The model was fitted in python using *l*_2_ logistic regression in the Scikit-learn package^90^ with the following steps for data collection and cross-validation. The time series data was sampled randomly around spike events, with a balance of *P*(*spikes at time t*) ≃ 0.5. We determined strength of *l*_2_-form regularization using 5-fold cross-validation. In each fold, we held 20% of the data for validation (completely non-overlapped with the rest of the data in the original full time series) and used the remaining 80% of the data for training. Moreover, we used square root form of the swim vigor data in fitting. We used model without regularization on validation data as the full model and determined the performance, using negative log likelihood (NLL), as

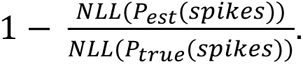

This is equivalent to a measure of the explained variance in a logistic regression.

Furthermore, we assumed the change of subthreshold membrane potentials linearly as

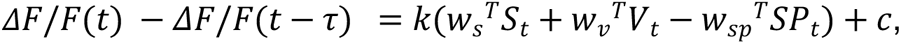

and thus the model error is

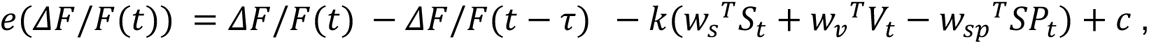

where k and c are two scalars in model fits.

We then fit the combined model of spike events and the change of subthreshold membrane potential as

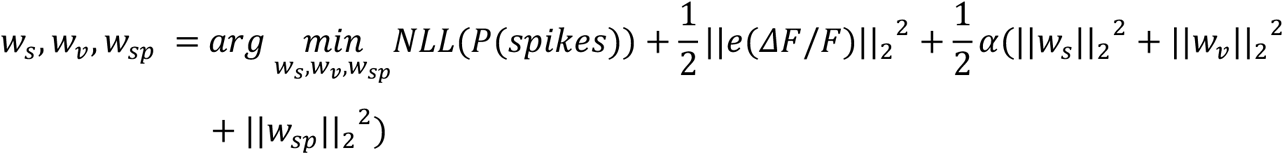

where *α* is *l*_2_-form regularization of the kernels. We performed graduate descent-based minimization with the initialization of the parameters fitted from the spike-event-only model above.

### Neural population codes

We computed the population coding of motor vigor learning by serotonergic neurons (**Fig. 6f, S7e**) using a sparse version of linear discriminant analysis^91^:

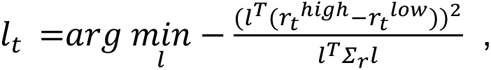

where a *l*_2_ regularizer, *γ* ∈ [0, 1], was applied to covariance matrix of the sample data

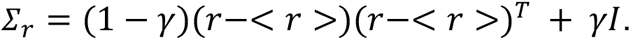

We sampled the data from the late phase of each G_ms_ period. The sample window for each data point to compute firing rate is 1 second.

### Spiking neural network model

We simulated the neural network dynamics in the dorsal raphe nucleus (**Fig. 5**) by using a spiking neural network model. Our model was equipped with 400 serotonergic neurons and 100 GABAergic neurons with sparse random connection at p = 0.03. The serotonergic neurons are modeled as adaptive integrate-and-fire neurons^92^:

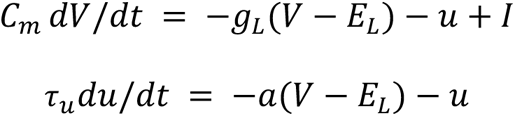

V is the membrane potential; u is the adaptation variable; I is the input current; *C*_*m*_ = 0.5 nF is the membrane capacitance; *g*_*L*_ = 0.025 uS is the leak conductance; *E*_*L*_ = −60 mV is the leak reversal potential; *a* = 0.01 uS is the adaptation coupling parameter (presenting the factor of the excitatory rebound currents) and *r*_*u*_ = 200 ms is the adaptation time constant for rebound currents). A spike happens when *V* > *V*_*t*L_ = −40 mV and V is then reset to -50 mV for a 2-ms refectory period; at the same time, spike triggers an inhibitory afterhyperpolarization, which would add onto adaptation variable,

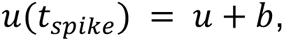

and b=0.001 nA. GABAergic neurons are modeled as simple integrate-and-fire neurons:

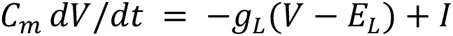

with nF; *g*_*L*_ = 0.02 uS; *E*_*L*_ = −60 mV.

The synaptic inputs are modeled as

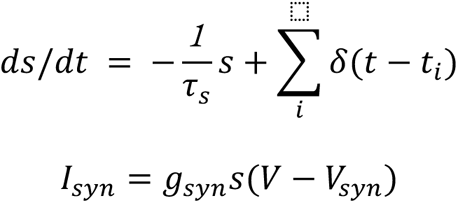

where *g*_*syn*_ is a synaptic conductance, *V*_*syn*_is the synaptic reversal potential, and s is a synaptic gating variable (that increases by one at a presynaptic spike event then decays at time constant *r*_*s*_). We set *r*_*s*_ = *50* ms; *g_5_*_–*HT*_ = *0*.*13* ns; *V_5_*_–*HT*_ = *0* mV; *g*_*GABA*_ = *0*.*52* ns; *V*_*GABA*_ = −*70* mV.

We provided excitatory and inhibitory inputs to serotonergic and GABAergic during the motor vigor learning task based on the results of neurotransmitter imaging. Both swim vigor and visual inputs were modeled as exponential decays from the behavioral variables onset times:

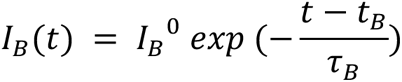

where the decay time constants were at 50 ms; max input *I*_*B*_^0^ = 100 pA for swim vigor; *I*_*B*_^0^ = 50 pA for visual inputs. Total inputs to a serotonergic neuron was thus

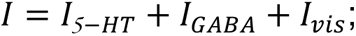

The total input to a GABAergic neuron was thus

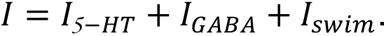

### Two-photon, plasma ablation of DRN GABAergic neurons

We performed two-photon plasma ablation of DRN GABAergic neurons and voltage imaging of serotonergic neurons before and after the ablation (**Fig. 6, S7**) by using triple transgenic zebrafish *Tg(tph2:Gal4; UAS:Voltron; gad1b:loxP-RFP-loxP-GFP)* and an optical setup described in our previous work (Vladimirov *et al.,* 2018). Briefly, we first performed a volumetric scan of red fluorescence, which represents GABAergic neurons expressing RFP, using 561-nm CW laser (Omicron, Germany) and 630/92 nm filter (FF01-630/92, Semrock), to determine the site of ablation manually. Based on this acquired stack, we chose 40-70 GABAergic cells inside the DRN for the main ablation (**Fig. 6**) or outside the DRN for control ablation (**Fig. S7c-f**). The distribution of selected points are shown in **Fig. S7a and b**. After the selection, we tuned the wavelength of a femtosecond laser (Camereon Ultra, Coherent) to 1050 nm and performed automatic ablation of selected cells. The lateral and depth positioning of the laser was controlled by a 2-axis galvanometer (Cambridge technology) and piezoelectric drive (Physik Instrument) of the objective lens. We chose dwell time per cell to 10-15 millisecond because of the depth of this area. There are >100 GABAergic neurons in the DRN (Kawashima et al., 2016), and we did not target all of them to avoid the risk of collateral damage to nearby serotonergic neurons.

We performed voltage imaging of serotonergic neurons that express Voltron conjugated with JF525 fluorescent dye for 5 minutes during motor vigor learning task before and after the above ablation procedure.

## Supplementary Figures

**Figure S1:**
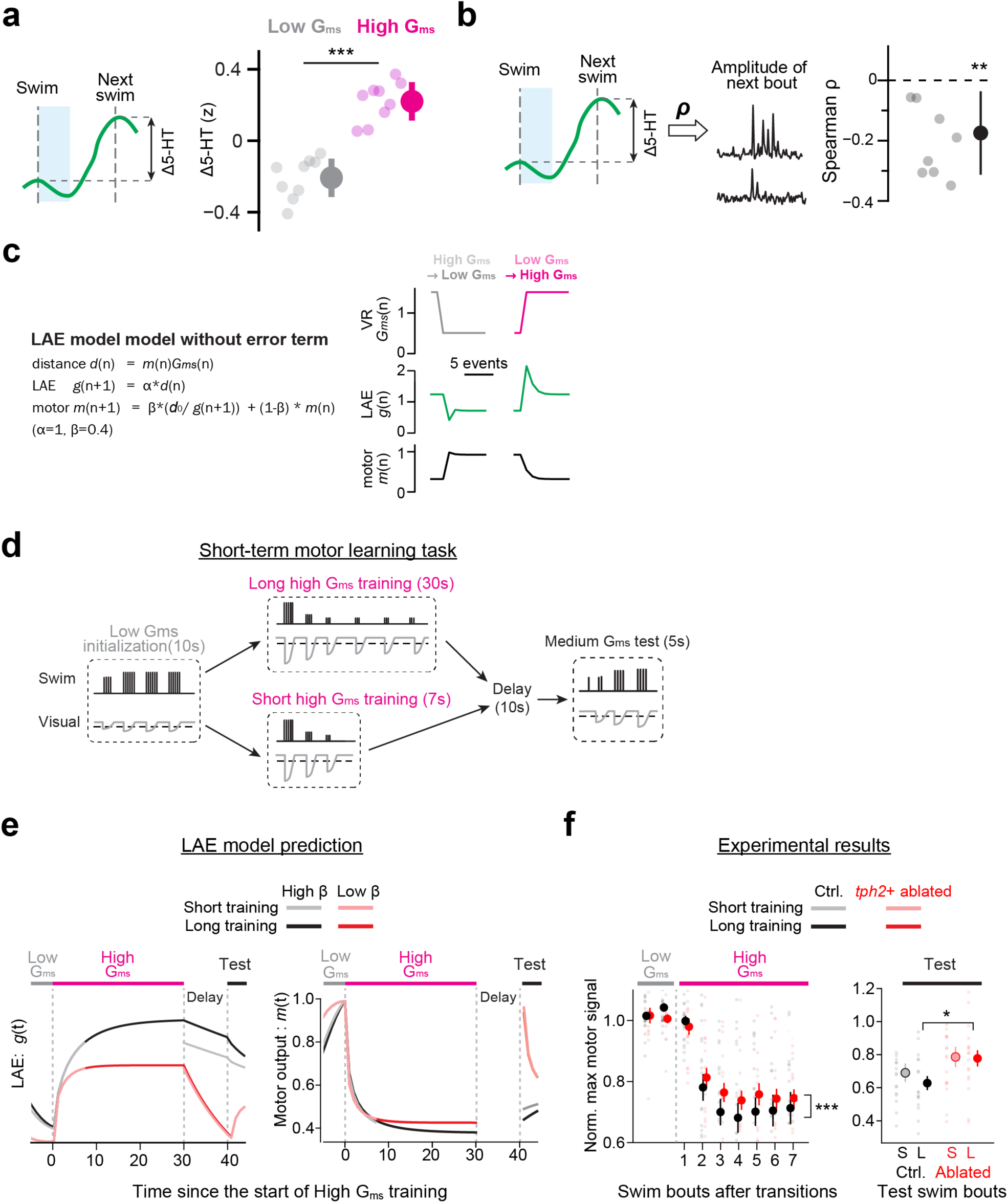
Serotonin release reflects learned effectiveness of action and modulates motor vigor learning. **(a)** The post-swim increase of serotonin in the hindbrain ROI (Fig. 1e), Δ5-HT, is high and positive in high Gms (magenta) and low and negative in low Gms (gray); N = 9 fish. ***, p=1.1*10^-5^, two-tailed paired signed-rank test. **(b)** The change in serotonin levels, Δ5-HT is negatively correlated with vigor of the next swim bout. Higher Δ5-HT corresponds to less vigorous subsequent swim bout; N = 9 fish. **, p = .0073, two tailed signed-rank test against zero. **(c)** Simulation of an alternative model for learning action effectiveness that does not involve error computation. *Left*, formula of the alternative learned action effectiveness (LAE) model. *Right*, the result of the simulation for 10 swim episodes at the transition from low G_ms_ to high G_ms_, and the transition from high G_ms_ to low G_ms_ during motor vigor learning. The temporal patterns of learned action effectiveness after changes in G_ms_ (middle) are significantly different from experimentally measured serotonin dynamics in the hindbrain of zebrafish (Fig. 1d). **(d)** Short-term motor learning paradigm for larval zebrafish. This behavioral paradigm from our previous work (Kawashima et al., 2016) tests not only the effects of changes in G_ms_ on swim vigor but also persistent behavioral effects of learned action effectiveness on swim vigor after a delay period. Swim vigor during the test period depends on the duration of high Gms training before the delay period. **(e)** Simulation of LAE model with error computation (Fig. 1f) for short-term motor learning paradigm in (d). Swimming occurs at 1 Hz. We simulated cases when the retention factor of the LAE model, β, was either high (0.98, black traces) or low (0.9, red traces). High β resulted in slow ramping of learned action effectiveness during the training and its slow decay during the delay period, which is qualitatively similar to the activity of raphe serotonergic neurons observed in our previous study. High β also resulted in [i] more suppression of swim vigor during the training and [ii] differences of swim vigor during the test period depending on the duration of the training. Duration of the delay period was simulated as 10 swim events, equivalent to about 10 seconds in real fish. **(f)** Experimentally observed changes in swim vigor during short-term motor learning paradigm in control fish (black) and fish whose *tph2+* raphe serotonergic neurons are ablated (red). The effects on swim vigor by the ablation are qualitatively similar to the simulation of LAE model with low β value in (e). N=17 and 16 for control and ablated fish, respectively. ***, p=6.4*10^-38^ for the difference between the control and ablated groups by two-way ANOVA test with repeated measures. *, p=0.034 for the difference of test swim vigor after long training between the control and ablated groups by two-sample t-test. This is a re-analysis of published data from Kawashima *et al*., 2016.

**Figure S2:**
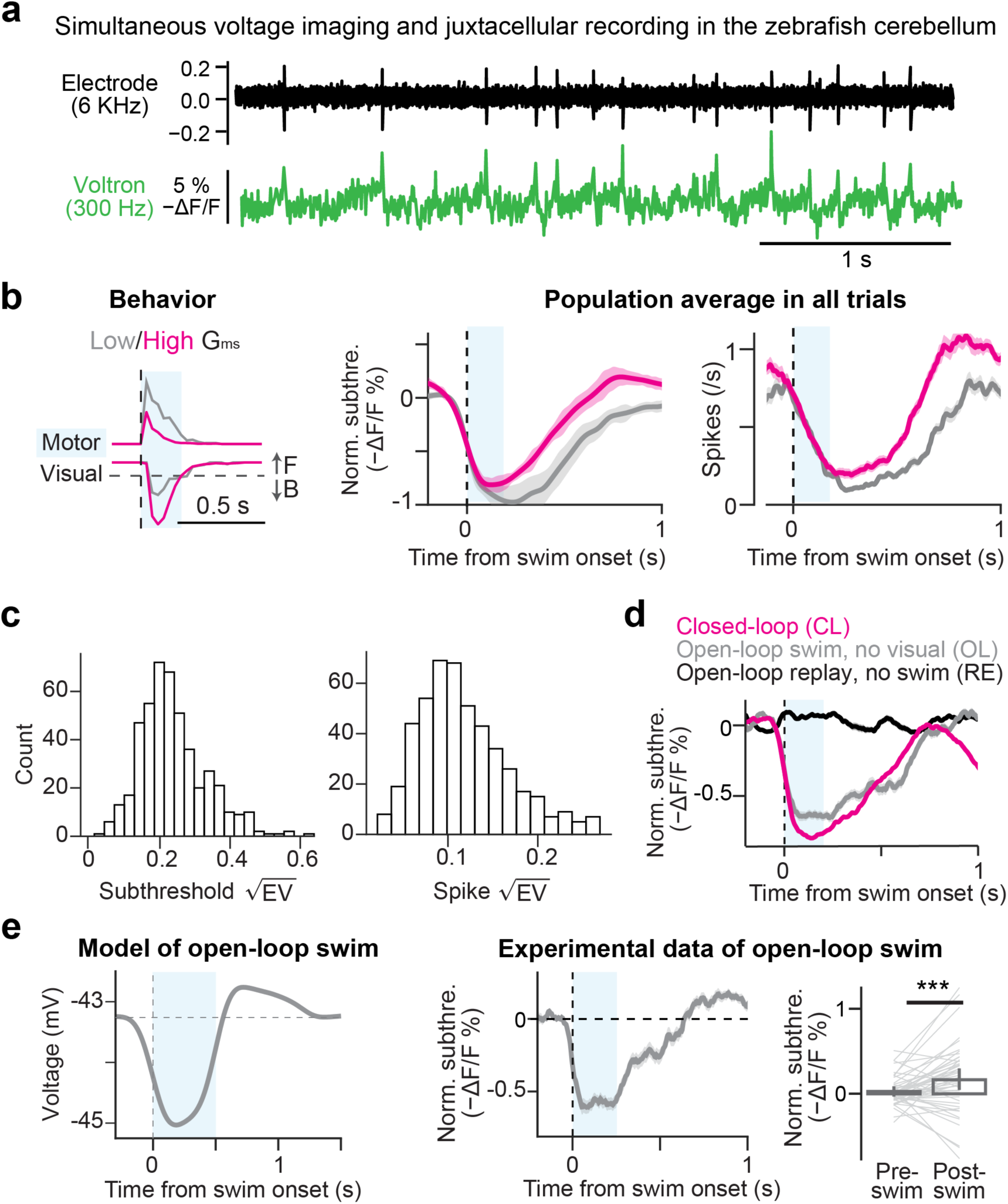
Supporting materials of spiking and voltage dynamics. **(a)** Signal traces of simultaneous voltage imaging and electrophysiology in eurydendroid cells in the zebrafish cerebellum, which were used for developing neural networks for spike detection in Fig. 2c. This data was acquired in Abdelfattah et al., 2019 ^30^. **(b)** Swim-triggered average of behavioral traces, membrane potential and spiking dynamics in serotonergic neurons during motor vigor learning. Compared to the motor-equalized analysis presented in Fig. 3d, here we present the triggered averages of all swim events. **(c)** Distribution of explained variance for subthreshold membrane potential and spiking activity for the model fitting shown in Fig. 3e. **(d)** Swim-triggered (magenta, gray) or visual-triggered (black) averages of membrane potential dynamics for closed-loop or open-loop experiments presented in Fig. 3f. We subsampled swim events to equalize swim vigor between closed-loop and swim only events for this analysis. **(e)** Rebound excitation during open-loop events in the serotonergic neurons. Simulation (left, see Fig. 5 for details) and experimental data (right) are presented. We subsampled open-loop swim events with strong vigor from the dataset of Fig. 3f and S3c to investigate the effects of rebound excitation after motor-evoked inhibition. N = 7 fish, 51 high G_ms_ cells. **\*\*\*,** p < .0001, paired signed-rank test in comparison of pre-versus post-swim membrane potential.

**Figure S3.**
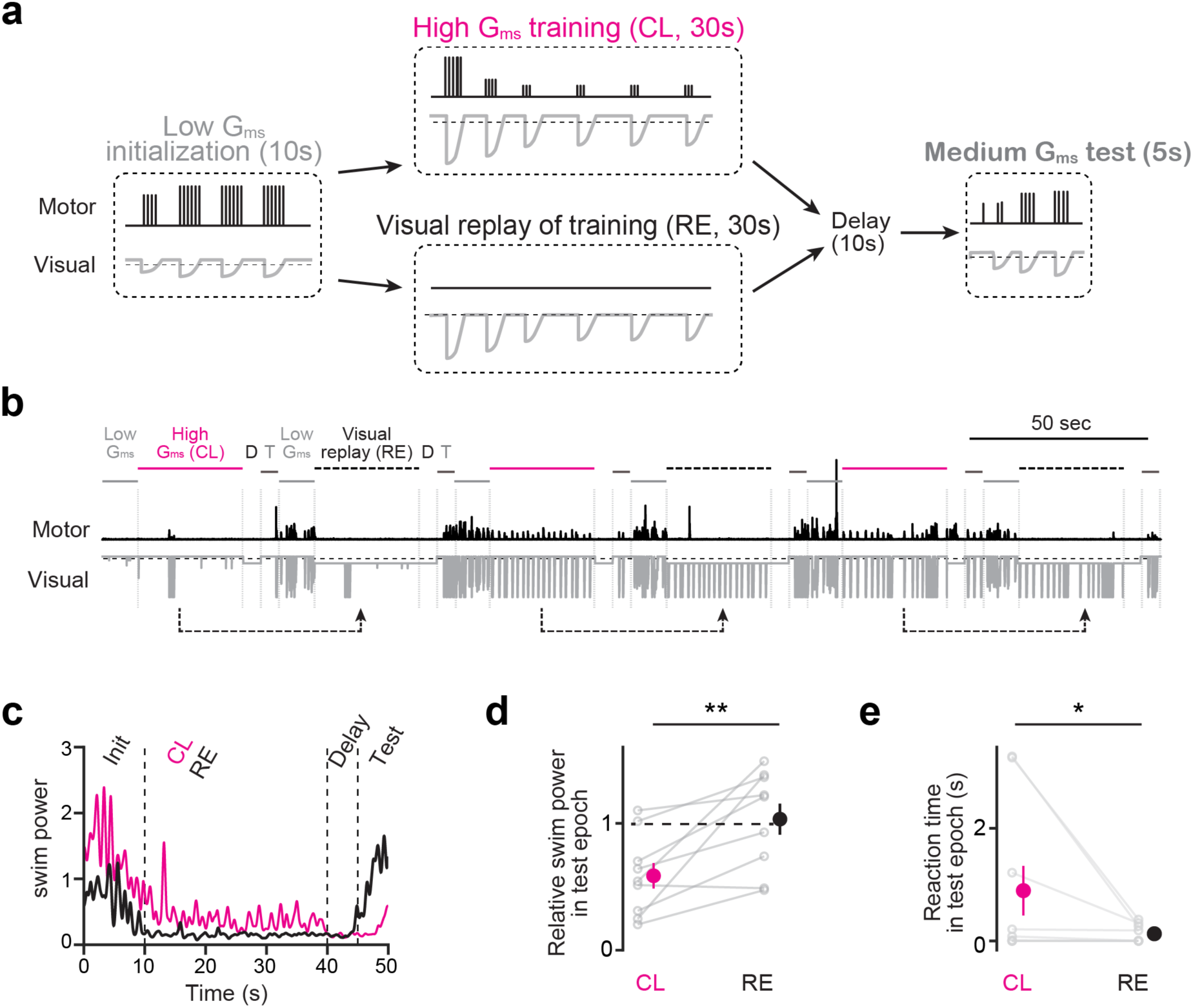
Less learning effects from swim-decoupled visual feedback during short-term motor learning paradigm. **(a)** Motor learning task that compares the learning effects of swim-coupled, closed-loop visual feedback (CL, top) and swim-decoupled, replayed visual feedback (RE, bottom). To prevent the occurrences in coincident swimming, we only replayed the backward component of the visual motion during the replay period. **(b)** Task structure (top), swimming patterns (middle) and visual stimuli motions (bottom) in 3 consecutive trials of the behavioral paradigm presented in (a). Of note, the fish swim very little during the replay in our analysis. **(c)** Average swim pattern during high G_ms_ closed-loop training period (magenta) and replay period (black) of an example fish. **(d)** Swim vigor decreases after high G_ms_ training but remains unchanged after replay. N = 9 fish. **, p = .0078, paired signed-rank test. **(e)** Reaction time becomes longer after G_ms_. N = 9 fish. *, p = .028, two-tailed paired signed-rank test.

**Figure S4:**
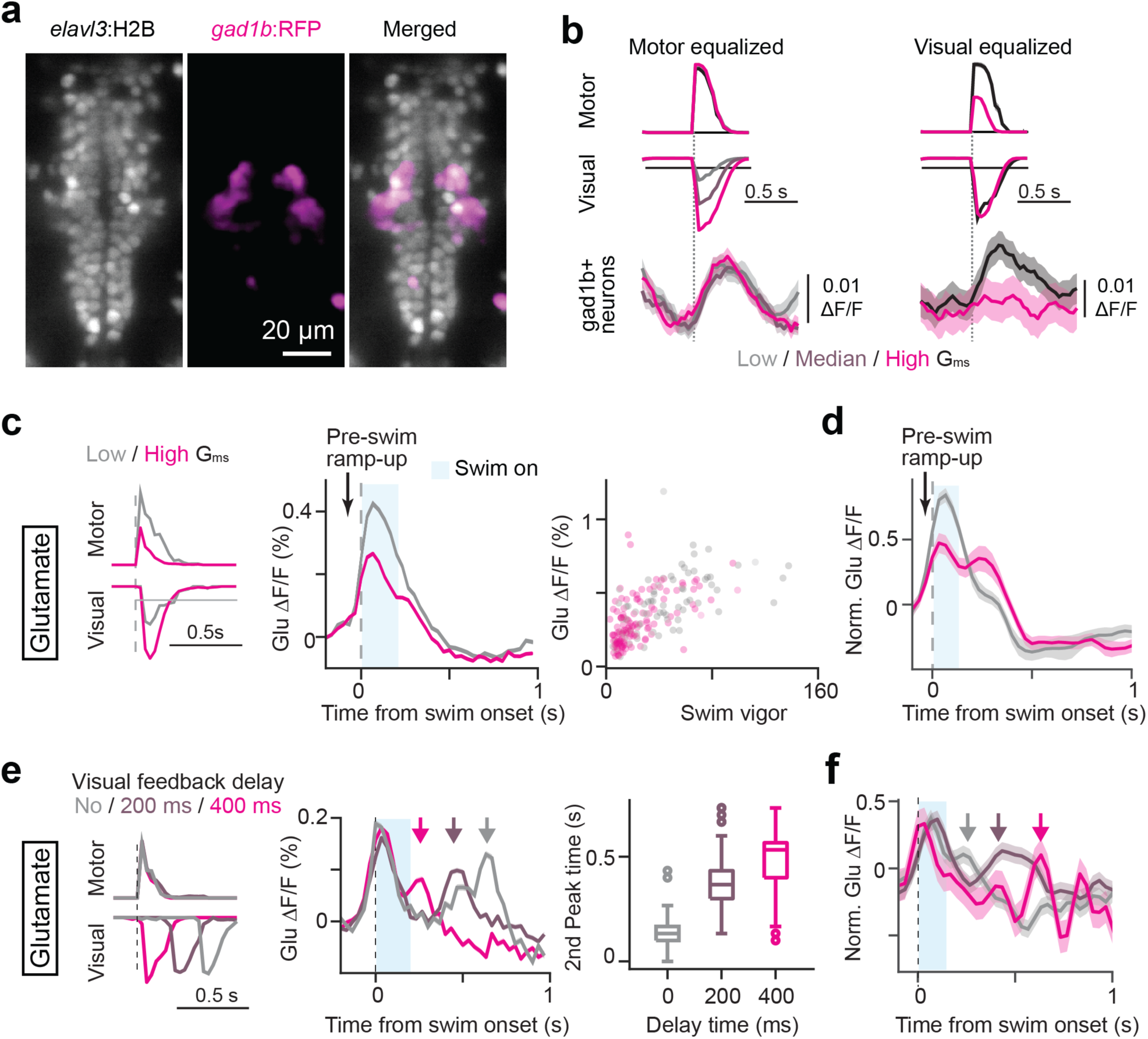
Supporting data for neurotransmitter imaging in the dendrites of serotonergic neurons. **(a)** Anatomy of raphe GABAergic neurons in the raphe nucleus that express red fluorescent protein (RFP) in the downstream of *gad1b* promoter in a transgenic zebrafish. **(b)** Raphe GABAergic neurons typically encode swim vigor rather than the velocity of visual feedback. (a, b) are replots of data collected in our previous study (Kawashima *et al.* 2016) ^14^. **(c,d)** Swim-triggered averages of glutamate inputs into the dendrites of raphe serotonergic neurons in an example fish (c) and across fish (d) during motor vigor learning paradigm. Unlike the analysis presented in Fig. 4e, we did not subsample swimming events to equalize swim vigor across different G_ms_ conditions in this analysis. N=17 fish for cross-fish analysis in (d). Shaded regions represent s.e.m. across fish. **(e)** Temporal shifts of glutamate inputs during delayed visual feedback of swimming. Example from a single fish. We equalized the vigor of swim bouts across conditions of a variable delay of visual feedback randomly given at 0ms, 200 ms, or 400 ms. *Right*, the second peak of the glutamate inputs shifted according to the set delays accordingly. **(f)** Cross-fish average of glutamate inputs in response to temporal delays of visual feedback after swimming shown in (e). N = 2 fish, 6 sessions. Shaded regions represent s.e.m. across sessions.

**Figure S5:**
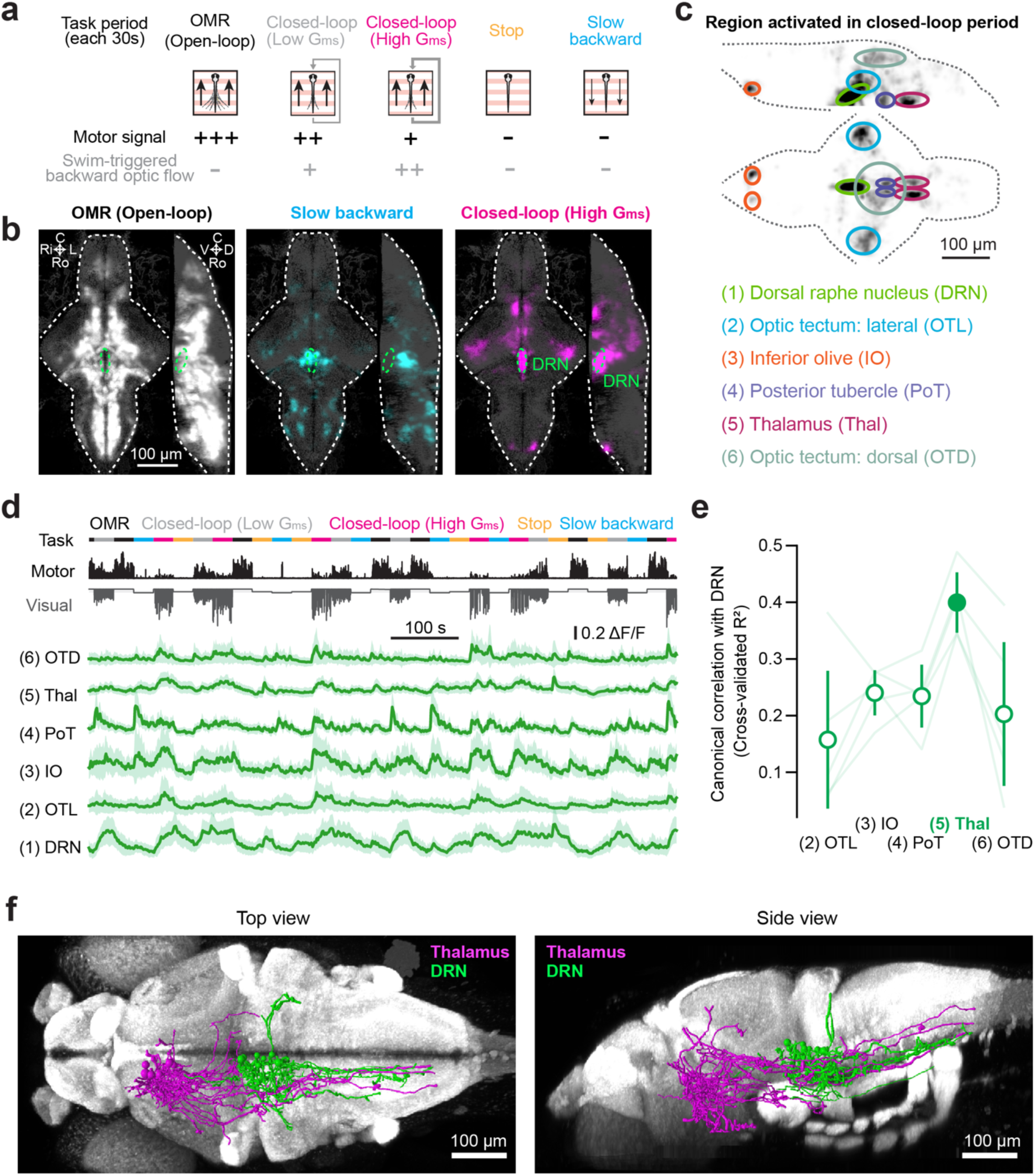
Potential upstream brain area that send visual information to raphe. **(a)** The behavioral task for identifying neurons activated by closed-loop sensory feedback. In the OMR (open-loop) period, the visual environment moves forward without any feedback. In the ‘Closed-loop’ period, the visual environment moves forward, but when a swim signal is detected it moves backward with low (gray) or high (magenta) G_ms_. In the ‘Stop’ or “Slow backward” period, the visual environment stops or slowly moves backward, during which the fish tend to swim less or cease swimming. **(b)** Whole-brain maps of neurons that showed highest activity in the OMR (open-loop) period (white), in the Slow backward period (cyan), or in the closed-loop high G_ms_ period (magenta) morphed and overlaid on a reference brain (gray). N = 5 fish. See Methods and Resources for details of analyses. D, dorsal; V, ventral; Ro, rostral; C, caudal; Ri, right; L, left. **(c)** Anatomical segmentation of brain regions that showed highest activity in the closed-loop G_ms_ period. *Top*, the same whole-brain map in (b) inverted for brightness and overlaid with anatomical masks. *Bottom*, a list of six identified anatomical regions. Brain areas showing G_ms_ encoding include: dorsal raphe nucleus (DRN), lateral optic tectum (OTL), dorsal optic tectum (OTD), posterior tubercle (PoT), thalamus (Thal) and inferior olive (IO). (a)-(c) are re-adapted from the supplementary figure of Kawashima *et al*., 2016 ^14^. **(d)** Population activity dynamics of six brain areas identified in (c). We extracted neurons that show the highest activity during the closed-loop (high G_ms_) period from each area and plotted their average activities from a representative fish. **(e)** Canonical correlation analysis between population dynamics of brain areas identified the thalamus as a potential upstream area of the DRN. For each fish, we extracted neurons that show the highest activity during the closed-loop (high G_ms_) period from each area and calculated cross-validated canonical correlation (R^2^) to neural populations in the DRN. We then further averaged these values across 5 fish. Error bars represent s.e.m. across fish. **(f)** Single-cell projection patterns from the thalamus (magenta) spatially overlaps with the dendrites of raphe serotonergic neurons (green). We used the single-cell projection atlas from Kunst et al., 2019 ^42^.

**Figure S6:**
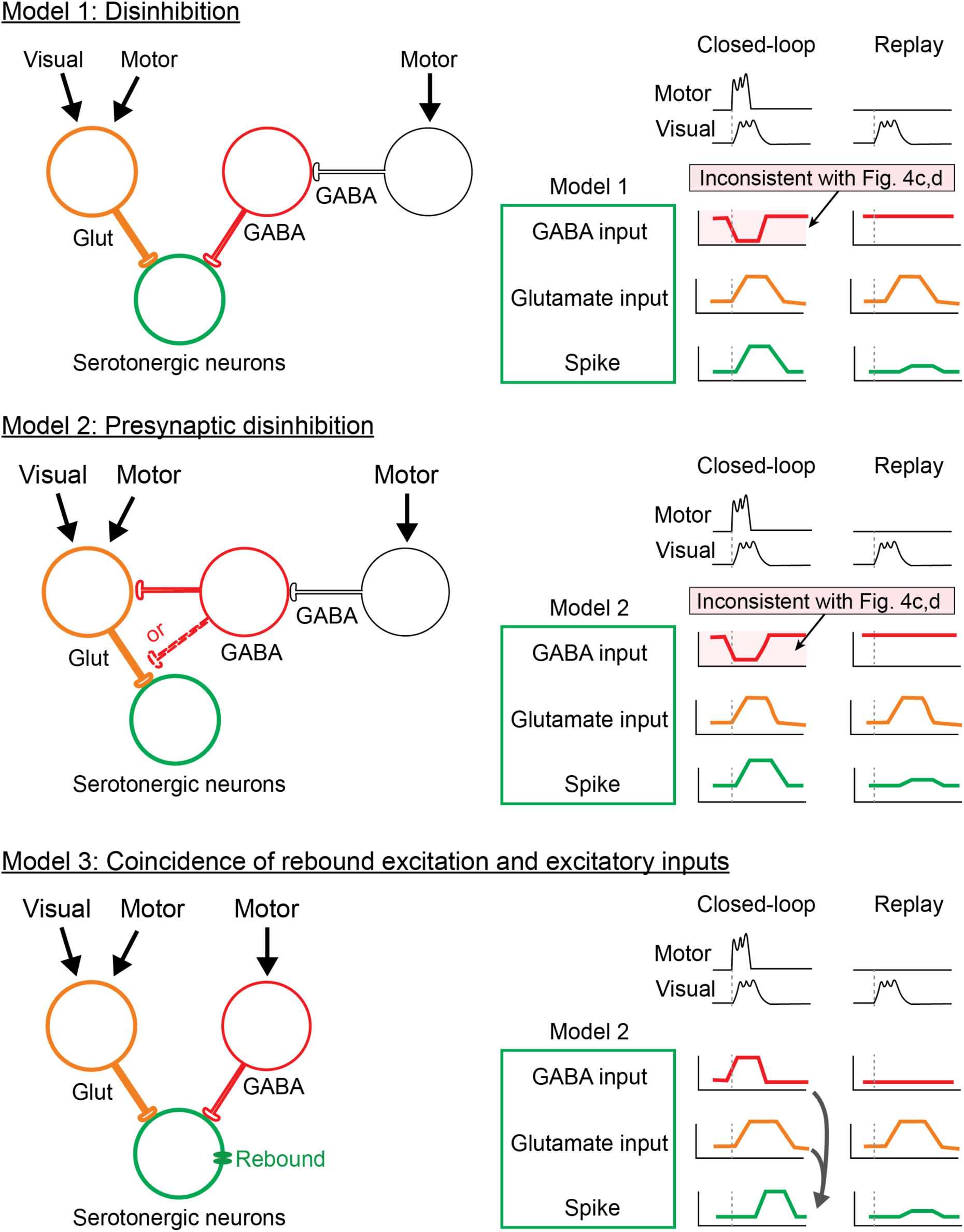
Candidate circuit models for sensory-motor computations in serotonergic neurons. Hypothetical circuit models of motor-gated sensory responses in raphe serotonergic neurons. Note, these model schematics do not represent an exhaustive list of possibilities. *Left*, we formulated models based on somatic disinhibition (Model 1), presynaptic disinhibition (Model 2) and coincidence of rebound excitation and synatic inputs (Model 3). *Right,* schematics of expected input patterns of GABA and glutamate. Only Model 3 is consistent with our experimental observation of GABA and glutamate inputs in Fig. 4.

**Figure S7.**
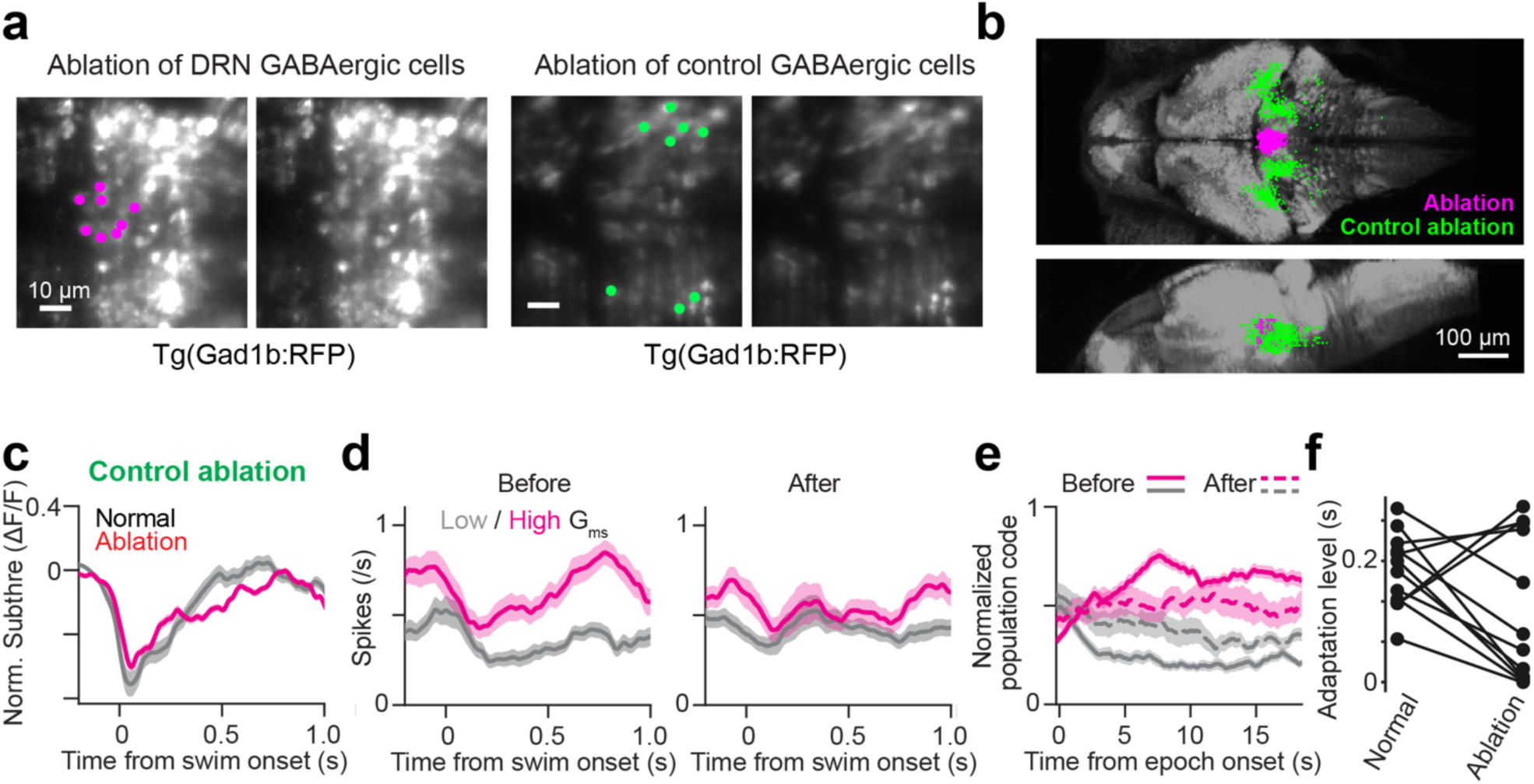
Supplementary materials for ablation of GABAergic neurons in DRN. **(a-b)** Anatomical loci of the ablation of raphe GABAergic neurons (magenta), presented in Fig. 6, and the control ablation of GABAergic neurons outside the raphe nucleus (green), presented in c-f. (a) Representative loci of two-photon plasma ablation in individual fish. (b) Summary of ablation loci for N=8 and 10 fish for the ablation of raphe GABAergic neurons (magenta) and GABAergic neurons outside the raphe (green), respectively. **(c)** Changes of membrane potential dynamics in serotonergic neurons before and after the control ablation. We did not see major reduction of swim-evoked inhibition after the control ablation compared to what we observed after the ablation of raphe GABAergic neurons in Fig. 6d. N = 9 fish, 48 high G_ms_ cells. Shaded regions represent s.e.m. across cells. **(d)** The control ablation of GABAergic neurons outside the raphe nucleus resulted in only partial loss of visual feedback encoding after swimming. *Left*, spiking in serotonergic neurons before the control ablation during low (gray) and high (magenta) G_ms_. *Right*, spiking in 5-HT neurons in the same animals after the control ablation. N = 9 fish, 48 high G_ms_ cells. Shaded regions represent s.e.m. across cells. **(e)** DRN serotonergic neurons retain population coding of *L*AE after the control ablation over swim events. N = 10 fish, 20 sessions. Shaded regions represent s.e.m. across sessions. **(f)** The control ablation of GABAergic neurons outside the raphe nucleus did not cause a reduction in motor adaptation. N = 12 fish, p = .129, paired signed-rank test in comparison of before and after the ablation.

